# Mosaic evolution of avian brain compartments revealed by comparative MRI

**DOI:** 10.64898/2026.07.01.735787

**Authors:** Yuki Kawabe, Tomokazu Tsurugizawa, Chiaki Ohtaka-Maruyama, Takuma Kumamoto

**Author notes:** Corresponding author: Chiaki Ohtaka-Maruyama and Takuma Kumamoto. **Author Contributions:** T.K. designed the research project. Y.K. analyzed MRI images. T.T. performed MRI analyses. T.K. wrote the article draft. C.O-M. supervised the study and revised the manuscript. T.K., T.T., and C.O-M. acquired funding. All authors revised the draft manuscript and approved the final version of the manuscript. **Competing Interest Statement:** The authors declare that they have no conflicts of interest to disclose. **Classification:** Biological Sciences, Evolution.

## Abstract

Birds evolved large, cognitively capable forebrains independently of mammals, yet comparative analyses of avian brain organization have been constrained by the lack of standardized resources capable of resolving internal parcellation and long-range connectivity across species. Here, we present a comparative MRI resource spanning 16 avian species representing major clades and diverse ecological niches. We analyzed high-resolution T2-weighted and diffusion-weighted datasets suitable for direct interspecific comparison. T2-weighted morphometry revealed pronounced region-specific variation in internal brain architecture, including lineage-dependent differences in the relative prominence of major brain divisions and commissural structures, supporting a pattern of mosaic diversification rather than uniform scaling. To validate MRI-derived anatomical boundaries, we compared MRI parcellations with complementary histological analyses in three representative taxa (the large-billed crow, gentoo penguin, and mandarin duck), demonstrating close correspondence between MRI-defined borders and cytoarchitectonic transitions identified by Nissl staining, as well as major myelinated compartments visualized by Luxol Fast Blue staining. Moreover, diffusion MRI tractography and fractional anisotropy (FA) mapping further revealed both conserved and species-specific features of large-scale brain organization. Seed-based tractography of the optic lobe, dorsal cortex, cerebellum, and anterior cortex in chick, gentoo penguin, and large-billed crow revealed conserved within-compartment trajectory patterns alongside marked region-specific interspecific differences, particularly in optic-lobe-associated long-range trajectories. Whole-brain FA maps revealed complementary variation in regional microstructural organization across taxa. Together, this comparative MRI framework provides a cross-validated foundation for linking internal brain anatomy and long-range connectivity to ecological and evolutionary diversification in birds, with broader applications to comparative neuroanatomy across amniotes.

**Significance Statement:** How does the internal architecture of the brain reorganize as species adapt to diverse lifestyles? Traditional comparative neuroanatomy has largely relied on overall brain size and external morphology, providing limited insight into internal subdivisions and long-range connectivity. In this study, we present a standardized avian MRI resource and a reproducible analytical pipeline that enables direct cross-species comparisons of 3D morphometry and diffusion-derived connectivity across 16 bird species representing major clades and diverse ecological niches. MRI-derived anatomical boundaries are validated through complementary histological analyses, providing biological support for cross-species parcellation and white-matter measurements. Using this framework, we demonstrate that avian brains diversify in a strongly mosaic manner, with region-and pathway-specific changes that vary disproportionately across lineages rather than scaling uniformly with overall brain size. By moving beyond simple correlations between brain size and behavior, this resource provides a benchmark for testing hypotheses of mosaic brain evolution and for linking neural architecture to ecological and behavioral diversification across vertebrates.

## Introduction

The overall organization of the vertebrate brain is broadly conserved, yet only a few lineages have undergone dramatic and repeated expansion of specific brain divisions. The telencephalon is a prominent example, having expanded extensively in both mammals and birds and representing two independent evolutionary routes to large, cognitively capable forebrains (1, 2).

Mammals evolved a laminated six-layer neocortex, whereas birds achieved comparable behavioral sophistication with a pallium that is predominantly nuclear in its cytoarchitecture (1, 3). This architectural divergence, despite partial functional convergence, makes birds a powerful comparative system for identifying general principles underlying forebrain enlargement and specialization across vertebrates (1, 4). Birds also display substantial diversity in ecological niches, sensory systems, social behaviors, and cognitive capacities across lineages (5–9). Such variation is thought to reflect mosaic scaling of pallial and subpallial regions, together with differences in long-range white-matter organization and interhemispheric integration. However, comparative studies of avian brain evolution have relied primarily on neuron-count scaling analyses and CT-based endocast reconstructions (3,10). Although these approaches have provided important insights into overall brain size and external morphology, they cannot resolve internal brain parcellation or long-range tract architecture. Consequently, the field still lacks harmonized, publicly available datasets and analytical frameworks for cross-species comparisons of internal brain organization and connectivity. Existing avian MRI resources have largely been limited to species-specific atlases or diffusion tensor imaging (DTI) studies focused on song systems, rather than comparative whole-brain datasets spanning multiple avian lineages (9, 11, 12).

Here, we address this gap by assembling a comparative MRI resource encompassing 16 bird species representing major avian clades and diverse ecological niches. The dataset combines 13 species from the Animal Brain Collection (ABC) with three publicly available datasets and applies a unified framework for structural, morphometric, and diffusion MRI analyses (11,13–16). Building on the ABC framework, we integrate high-resolution structural and diffusion MRI data acquired using harmonized protocols. We register brains to species-specific templates and a common comparative space, enabling quantification of regional volumes and morphology, relative investment across major brain divisions, white-matter macroarchitecture, and commissural organization. Using this framework, we evaluate three a priori predictions regarding avian neuroanatomical diversification. First, associative pallial territories exhibit disproportionate scaling in taxa with complex social behaviors. Second, white-matter volume and tract coherence covary with relative telencephalon size. Third, interhemispheric routing displays clade-specific scaling consistent with alternative architectural solutions for bilateral integration. Together, this study provides a comparative MRI resource for birds and a reproducible framework for linking whole-brain anatomy and connectivity to ecological and behavioral diversification across avian lineages, with broader applicability to comparative neuroanatomy across vertebrates.

## Results

### MRI Analysis of a Comparative Structural MRI Resource Across Avian Taxa

First, we assembled a comparative avian MRI cohort comprising 12 species from the ABC (13) and four publicly available avian MRI datasets: canary (14), starling (16), pigeon (15), and zebra finch (11) (Fig. 1). The resulting dataset included 16 species representing four major avian groups: Galloanserae (green), Columbaves (purple), Aequorlitornithes (blue), and Inopinaves (orange). High-resolution T2-weighted and diffusion-weighted images were obtained for direct interspecific comparisons. Representative coronal sections demonstrated sufficient image quality and contrast to resolve internal anatomical landmarks across all taxa, providing a robust basis for subsequent morphometric and connectivity analyses (Fig. 1B).

**Figure 1.**
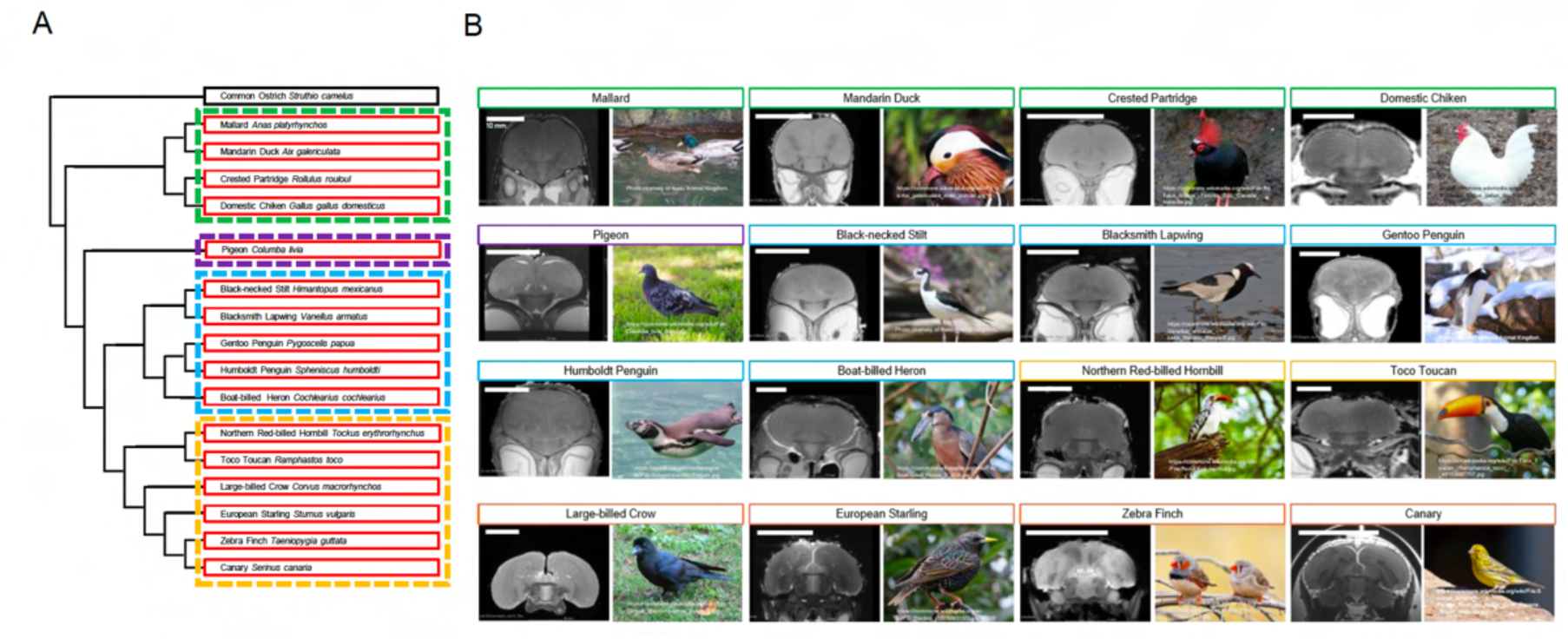
Avian species included in a comparative MRI dataset. Overview of the 16 avian species included in this study, representing major avian clades and diverse ecological niches. (A) Phylogenetic relationships among the sampled species, grouped into four major avian lineages. (B) Representative coronal T2-weighted MRI sections and corresponding photographs of each species. These images illustrate the diversity of brain sizes and overall morphologies represented in the dataset and demonstrate sufficient internal contrast to identify major anatomical landmarks across taxa. All MRI datasets were processed using a standardized preprocessing workflow to enable direct cross-species comparisons.

**Table 1.**
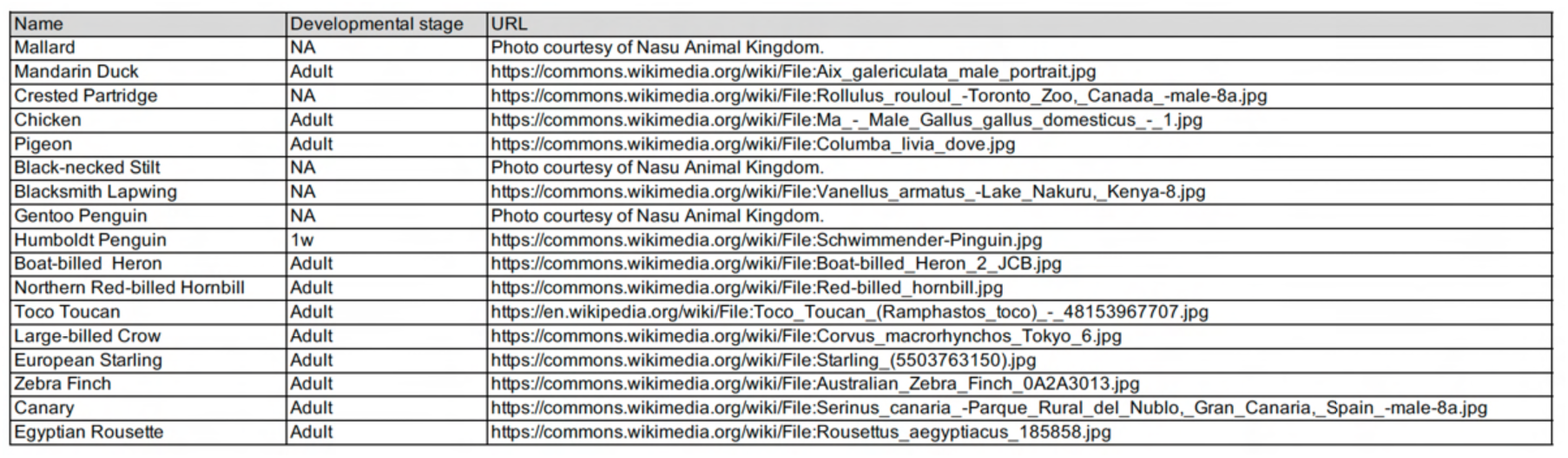
Image sources for animal photographs used in Figure 1.

### Interspecific Diversity in Internal Brain Architecture

T2-weighted morphometry revealed substantial interspecific variation in the relative size, morphology, and configuration of major brain divisions. For example, the anterior commissure (AC) was readily identifiable in all 16 species but differed markedly in its relative prominence among taxa (Fig. 2A). The AC appeared particularly prominent in the large-billed crow, starling, and canary, whereas it was comparatively reduced in the mallard, mandarin duck, and crested partridge. Penguins also exhibited a relatively small AC, suggesting lineage-dependent differences in telencephalic interhemispheric pathways across species (Fig. 2A). However, quantitative analysis showed that AC area scaled approximately with whole-brain area across species (Fig. 2B), indicating that relative AC size varies little after normalization to overall brain size. In birds, the AC interconnects arcopallial and preoptic/anterior forebrain regions and can also link non-homotopic territories between hemispheres (17). Although AC size cannot be interpreted directly as a measure of processing complexity, it represents the principal commissural pathway connecting the two pallial hemispheres in birds. Collectively, these findings suggest that the apparent interspecific differences in AC prominence are largely attributable to variation in overall brain size within the present dataset. Thus, while the AC remains an important anatomical landmark for comparative studies of avian interhemispheric organization, our area-based analysis provides limited evidence for major lineage-specific differences in relative commissural investment.

**Figure 2.**
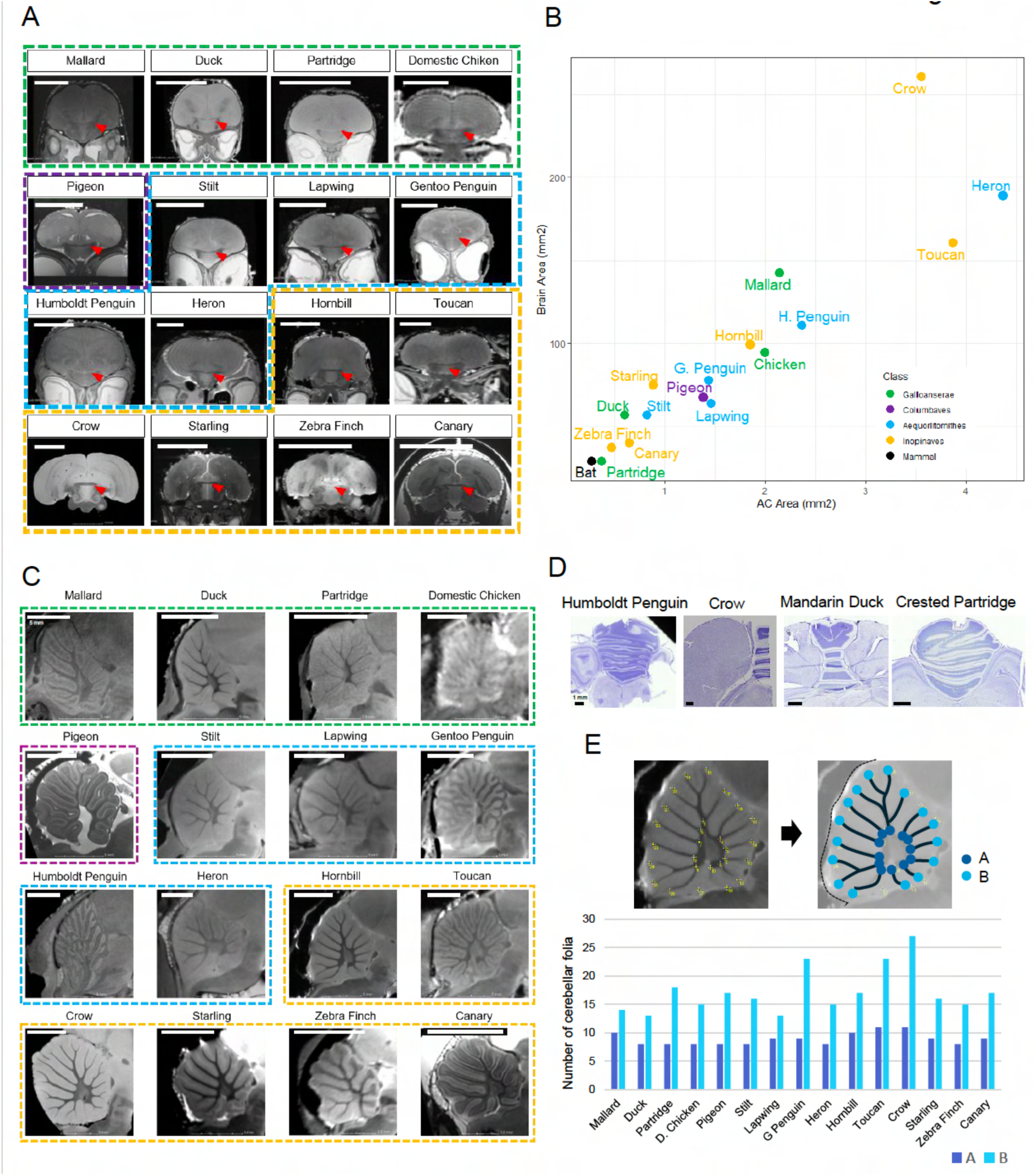
Whole-brain T2-weighted MRI highlights region-specific interspecific diversity in birds. (A) Representative coronal T2-weighted images from multiple species reveal pronounced differences in the relative size and morphology of major brain divisions, as well as overall head morphology. The anterior commissure (AC), the principal interhemispheric commissure in birds, is readily identifiable across taxa (red arrows). (B) Quantification of AC size relative to whole-brain area across species. AC area scales approximately with brain size, indicating that apparent differences in AC prominence largely reflect variation in overall brain size rather than substantial differences in relative commissural investment. (C) Representative sagittal T2-weighted images allowing qualitative comparison of cerebellar morphology across species. Cerebellar foliation and overall cerebellar expansion vary markedly among taxa, with particularly elaborate folial patterns observed in penguins, toucans, and crows. (D) Representative cerebellar histological sections from selected species illustrating interspecific differences in folial branching and structural complexity. (E) Quantification of cerebellar folial branching. The number of basal branches (A) exhibits relatively limited interspecific variation, whereas the number of terminal branches (B) varies more extensively and is elevated in species with more highly elaborated cerebella. Together, these findings support a pattern of mosaic, region-specific diversification in avian brain organization.

Next, we examined interspecific variation in cerebellar development using sagittal sections, which allow detailed assessment of cerebellar structural complexity (Fig. 2C). Sagittal T2-weighted sections revealed clear differences in the overall size and folial organization of the cerebellum among species. The mallard, mandarin duck, crested partridge, black-necked stilt, and boat-billed heron exhibited fewer and less elaborate cerebellar folia, whereas the pigeon, large-billed crow, and toucan possessed larger and more densely foliated cerebella. To further anchor these MRI-based observations to tissue-level anatomy, representative histological sections were examined in selected species and showed corresponding differences in folial branching patterns and cerebellar structural complexity (Fig. 2D). Quantification of cerebellar folial branching revealed relatively little variation in the number of basal branches across species, whereas terminal branch number was substantially greater in penguins, toucans, and crows (Fig. 2E). This distinction between basal and terminal branches indicates that interspecific variation was most evident in distal folial elaboration rather than in the primary branching scaffold of the cerebellum. This pattern suggests that the primary cerebellar scaffold is comparatively conserved across the sampled taxa, whereas distal folial elaboration represents a more variable anatomical feature. Because cerebellar foliation increases the available cortical surface within the cerebellum, variation in terminal branching may reflect lineage-specific expansion of sensorimotor processing capacity, although the present analysis does not directly resolve functional specialization. Notably, penguins displayed pronounced cerebellar expansion despite having a relatively small AC, suggesting a dissociation between the elaboration of interhemispheric and sensorimotor systems. Together, these observations are consistent with mosaic, region-specific diversification of internal brain architecture rather than uniform scaling across all major brain divisions.

### Template-Based 3D Morphometry and Mosaic Scaling

High-resolution MRI enables not only comparisons of overall brain morphology across species but also quantitative analyses of individual brain regions. Using the MRI datasets, we reconstructed three-dimensional (3D) brain models for 16 avian species and an Egyptian rousette bat, which was included as a flying mammalian outgroup. Representative renderings are shown from anterior-left (Fig. 3A, left) and posterior-right (Fig. 3A, right) perspectives. The quality and smoothness of the reconstructed 3D surfaces depended on the image acquisition resolution, with higher-resolution datasets producing more refined reconstructions. These renderings provide a clear visualization of the spatial relationships among major brain divisions, including the telencephalon, midbrain, and cerebellum, within each species.

**Figure 3.**
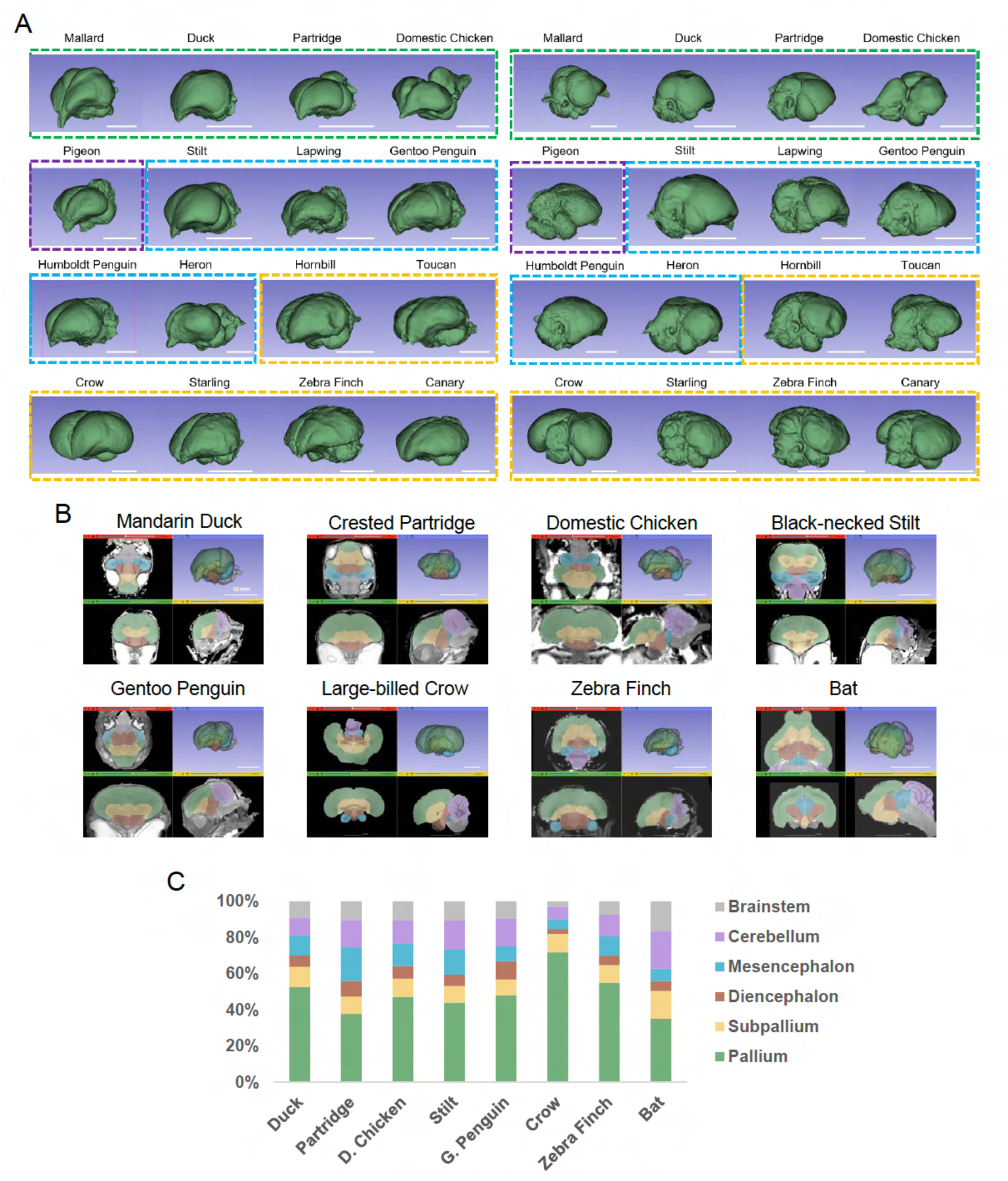
Template-based registration enables standardized 3D morphometry and mosaic scaling quantification. (A) Three-dimensional (3D) reconstructions of brains from 16 avian species, rendered from anterior-left (left) and posterior-right (right) viewpoints. The fidelity of the reconstructed surfaces reflects image acquisition resolution, with higher-resolution datasets producing smoother and more detailed reconstructions. (B) Schematic overview of the morphometric workflow, including species-specific template construction, registration into a common comparative space, and template-guided segmentation of major brain compartments. Representative overlays demonstrate the correspondence between MRI data and segmented 3D reconstructions. (C) Volumetric quantification of major brain divisions based on segmentation-derived voxel counts, expressed as proportions of total brain volume to facilitate comparison of relative regional investment across species. This workflow enables standardized evaluation of mosaic scaling across diverse avian lineages.

We next established a template-based morphometric workflow to convert these anatomical reconstructions into comparable regional measurements (Fig. 3B). In this workflow, each brain was aligned to a species-specific template, major brain compartments were segmented, and the resulting parcellations were used to generate 3D surface models and segmentation-derived volume estimates. This procedure allowed the same major anatomical compartments to be compared across species despite substantial differences in brain size and overall morphology. To quantify regional investment, we performed volumetric segmentation using species-specific templates and calculated the relative volume fractions of the telencephalon, midbrain, and cerebellum (Fig. 3C). The resulting proportional volume profiles suggested that overall brain size alone may not fully capture interspecific variation in internal brain organization. Instead, individual species showed distinct combinations of relative telencephalic, midbrain, and cerebellar allocation. For example, species with prominent cerebellar morphology did not necessarily show parallel increases in all other major brain divisions, suggesting that regional compartments can vary semi-independently. These 3D volumetric data therefore extend the two-dimensional observations from T2-weighted sections and provide an exploratory quantitative framework for evaluating mosaic scaling across avian lineages. This standardized workflow provides a foundation for phylogenetically informed analyses of mosaic scaling across diverse avian lineages.

### Histological Validation of MRI-Derived Boundaries

To assess the biological validity of the parcellation framework, we compared MRI-derived boundaries with complementary histological data from three representative species occupying distinct ecological and phylogenetic positions: the large-billed crow, gentoo penguin, and mandarin duck (Fig. 4). Across all three taxa, MRI-defined boundaries corresponded closely to histological transitions. Nissl staining confirmed that cytoarchitectonic changes aligned with the anatomical borders used for MRI segmentation, whereas Luxol Fast Blue (LFB) staining highlighted major white-matter compartments and long-range pathways identified by diffusion MRI. Together, these cross-modal correspondences provide tissue-level validation of the morphometric and diffusion-derived measurements and support the consistency of the parcellation framework across species.

**Figure 4.**
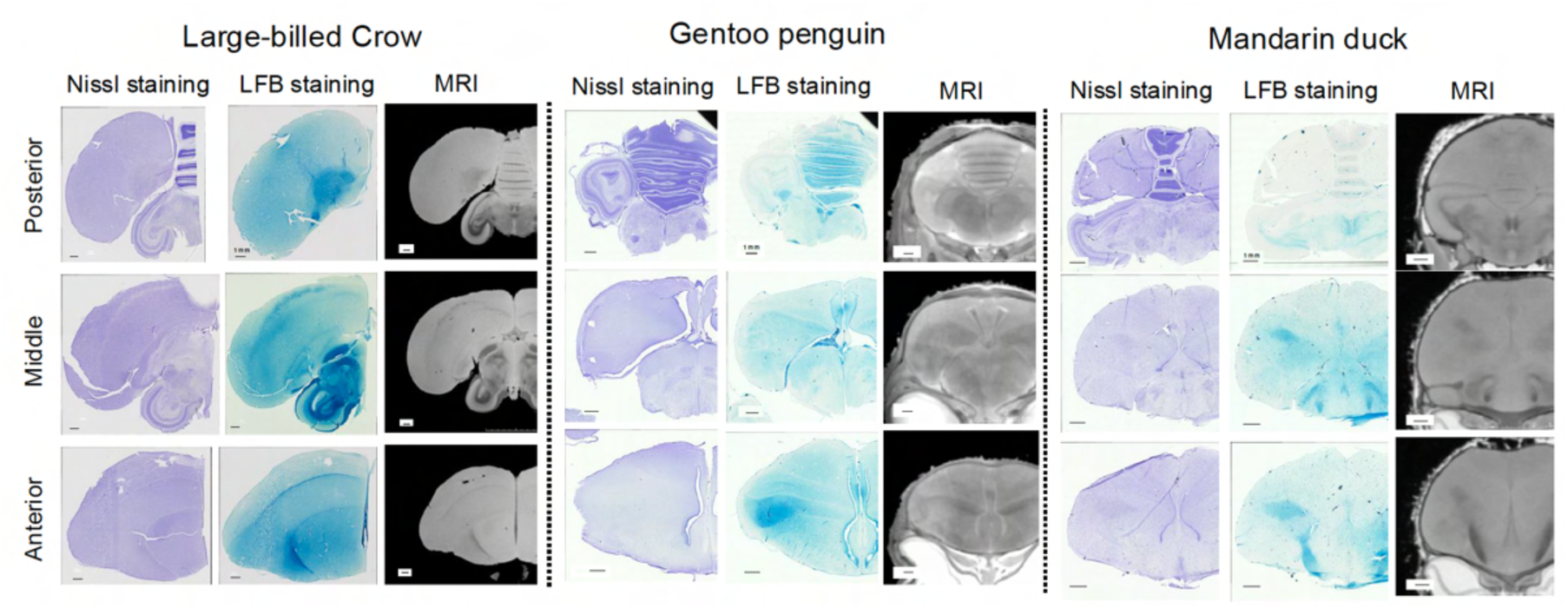
Histological validation links MRI-derived boundaries and diffusion-related measures to tissue architecture. (Left) Representative Nissl-stained sections across selected taxa illustrating cytoarchitectonic transitions that align with major MRI-defined anatomical boundaries used for parcellation. (Middle) Luxol Fast Blue (LFB) staining highlights white-matter architecture and major fiber-rich compartments corresponding to MRI-defined white matter. Together, these histological observations support the cross-species consistency of the segmentation framework and provide tissue-level validation for the interpretation of MRI- and diffusion-derived metrics.

### Divergent Long-Range Wiring and Microstructural Profiles

Diffusion MRI tractography and fractional anisotropy (FA) mapping revealed both conserved and species-specific features of avian brain organization (Fig. 5). We first examined region-seeded tractography to compare large-scale trajectory patterns among representative species (Fig. 5A). To compare long-range trajectory patterns, we performed tractography from four brain regions, namely, the optic lobe, dorsal cortex, cerebellum, and anterior cortex, in three representative species: chick, gentoo penguin, and large-billed crow. We reconstructed approximately 1.0 × 10^7^ streamlines and applied clustering analysis to identify the major trajectory groups. This analysis allowed us to compare how reconstructed trajectories associated with homologous or anatomically corresponding regions were distributed within each brain. The most pronounced interspecific differences were associated with optic-lobe tractography. In chick, reconstructed streamlines were largely restricted to the optic lobe, with only limited extension into lateral pallial territories. In contrast, optic-lobe-associated trajectories in the gentoo penguin were distributed more broadly throughout the brain. Pallial tractography also differed between species. In chick, dorsal-cortex-associated trajectories were dominated by dorsotemporal pathways and remained largely confined to pallial compartments. In the gentoo penguin, however, an additional distinct trajectory group was identified within a lateral temporal cortex-like subdivision. By comparison, cerebellum- and anterior-cortex-associated tractography showed broadly similar patterns in chick and the gentoo penguin, with many reconstructed trajectories remaining within their respective anatomical compartments. Thus, Fig. 5A indicates that some trajectory classes are broadly conserved across species, whereas optic-lobe- and pallium-associated trajectories show more conspicuous interspecific divergence. These findings indicate that tractography captures both conserved and region-specific patterns of organization across species.

**Figure 5.**
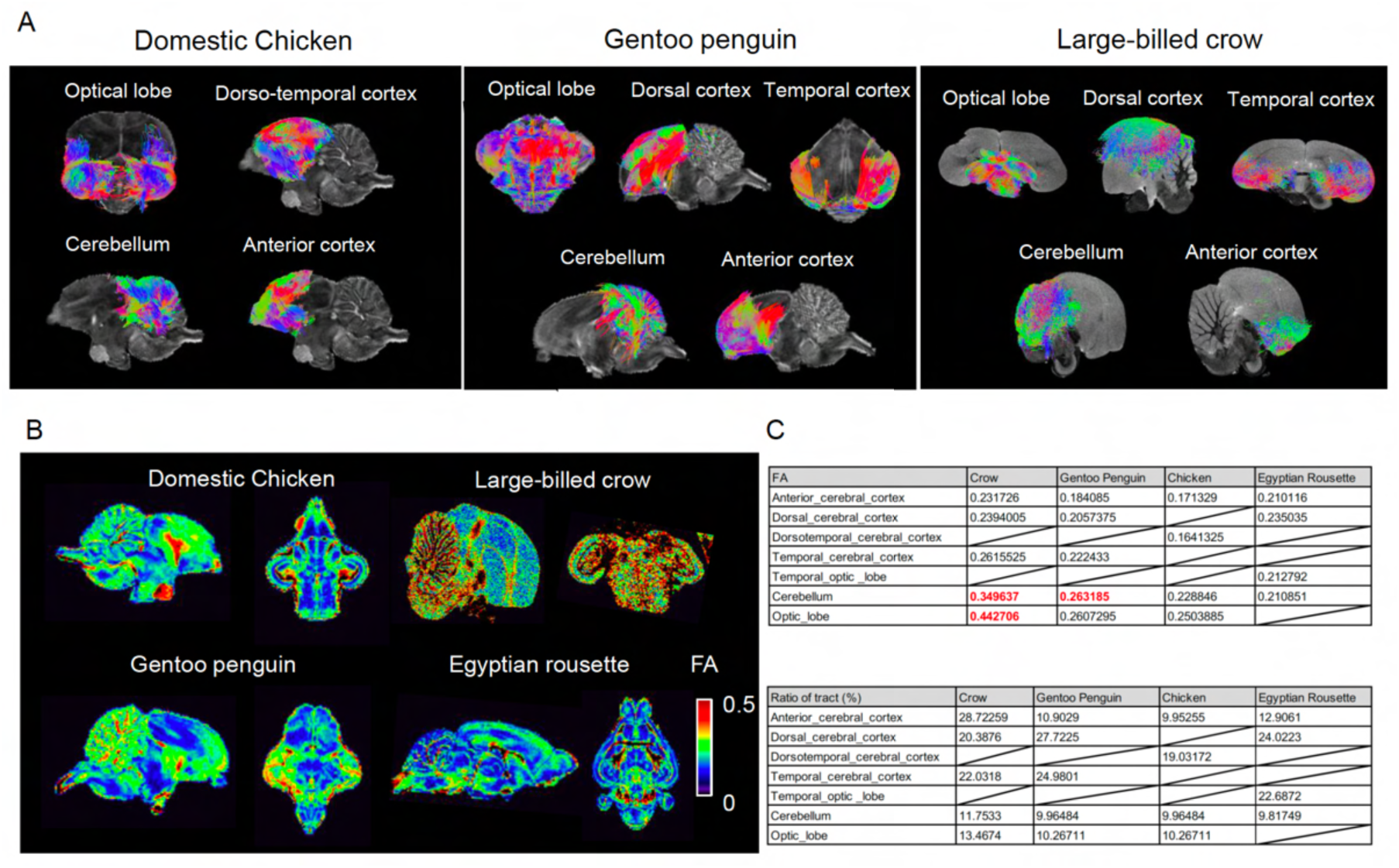
Diffusion MRI tractography and anisotropy mapping reveal conserved and divergent long-range organization across species. (A) Region-seeded diffusion MRI tractography in chick, gentoo penguin, and large-billed crow. Streamlines were generated from four regions of interest (ROIs)—the optic lobe, dorsal cortex, cerebellum, and anterior/temporal cortex—and subsequently clustered to identify the major trajectory groups. (B) Whole-brain fractional anisotropy (FA) maps from representative species illustrating interspecific differences in regional microstructural organization. (C) Quantitative comparison of tractography- and FA-derived regional metrics in chick, gentoo penguin, and Egyptian rousette bat. The upper and lower tables summarize relative regional distributions derived from reconstructed trajectories and FA values, respectively. Together, these analyses indicate that species share broad organizational features while differing substantially in inferred long-range routing patterns and regional microstructural profiles.

We next examined whole-brain FA maps to compare regional microstructural organization independently of streamline trajectory distributions (Fig. 5B, Movie 1-4). Whole-brain FA mapping further revealed interspecific differences in microstructural organization. In the large-billed crow, reconstructed trajectories were relatively enriched in the anterior lobe, whereas FA values were comparatively elevated in the cerebellum and optic lobe. In the gentoo penguin, FA values were likewise elevated in the cerebellum, although this was not accompanied by a corresponding increase in relative trajectory abundance. Instead, reconstructed trajectories were enriched in the dorsal cortex. In the mammalian outgroup, the Egyptian rousette, reconstructed trajectories showed a close association between the optic and temporal lobes, together with relative enrichment within the dorsal cortex. These observations suggest that FA-based microstructural contrast and tractography-derived trajectory abundance capture partially distinct aspects of brain organization. To summarize these relationships quantitatively, we compared regional values derived from reconstructed trajectories and FA maps across representative species (Fig. 5C). The tractography-derived summary highlighted species-specific differences in the relative distribution of reconstructed trajectories, whereas the FA-based summary emphasized regional variation in tissue anisotropy. The partial mismatch between these two measures was particularly evident in the gentoo penguin, in which cerebellar FA was elevated despite the absence of a corresponding enrichment of cerebellar trajectories. This dissociation indicates that regional microstructural specialization does not necessarily scale directly with the number or distribution of reconstructed streamlines.

Collectively, the tractography and FA analyses indicate that avian species share broad features of large-scale brain organization while exhibiting substantial differences in long-range trajectory distributions and regional microstructural profiles.

## Discussion

### A Unified Framework for Comparative Avian Neuroanatomy

Our study establishes a standardized MRI-based framework for quantifying internal brain organization across 16 avian species occupying diverse ecological niches. Previous comparative studies of avian brain evolution have relied primarily on measures of overall brain size, endocast morphology, or neuron-number scaling (3, 18–20). Although these approaches have yielded valuable insights, they cannot directly resolve internal anatomical organization or long-range pathway architecture. By integrating curated datasets from the ABC (13) with publicly available MRI resources, we provide a scalable and transferable platform for comparative neuroanatomy that can be extended to other vertebrate groups wherever structural and diffusion MRI data are available at the species level (21, 22). More broadly, this framework facilitates a shift from global brain-size correlations toward region-specific, phylogenetically informed hypotheses of neural diversification (18, 23).

### Mosaic Evolution and Ecological Diversification of Avian Brains

Our findings strongly support a mosaic model of avian brain evolution, in which individual brain divisions vary semi-independently rather than scaling uniformly as part of a single integrated structure (19). This interpretation extends previous work linking ecological variation to overall brain size by suggesting that ecological pressures are also reflected in the allocation of neural investment among internal brain regions (23–27). Under this framework, species facing distinct ecological challenges, such as rapid maneuvering, long-distance navigation, or underwater foraging, exhibit differential investment in sensory and sensorimotor systems, accompanied by corresponding changes in long-range white-matter organization (23, 28, 29).

Several observations from our dataset illustrate this pattern. The pronounced cerebellar expansion observed in the gentoo penguin, together with elevated FA within cerebellar white matter, is consistent with enhanced sensorimotor specialization associated with aquatic locomotion. By contrast, the AC exhibited conspicuous qualitative variation across species and appeared particularly prominent in large-billed crow, starling, and canary. However, because AC area scaled approximately with overall brain size in the present analysis, these observations should be interpreted with caution. Rather than providing evidence for lineage-specific expansion of commissural investment, they identify interhemispheric architecture as a potentially important axis of avian forebrain diversification that warrants further investigation using residual-based allometric approaches, broader taxonomic sampling, and tract-level analyses (23, 30).

More generally, variation in pallial and subpallial investment, cerebellar allocation, and the balance between commissural and association-type wiring patterns may represent complementary neural solutions to habitat-specific control demands that evolve semi-independently across avian lineages (18, 19). Brain evolution has traditionally been framed in terms of two major models, concerted evolution and mosaic evolution. An important unresolved question is when mosaic divergence emerges during development. Recent work in axolotl suggests that these mechanisms are not mutually exclusive but may jointly contribute to brain development and evolutionary diversification (31). Consistent with this possibility, post-hatching changes in avian brain morphology have been documented, and comparative studies indicate that sensory specializations, most notably in the olfactory bulbs and more broadly throughout major visual pathways including the optic tectum, are associated with ecological diversification in birds (5, 7, 32). Taken together, these findings raise the possibility that post-hatching brain development is further shaped in a species-specific manner by the ecological and behavioral demands experienced by individual lineages.

### Long-Range Wiring as a Dimension of Brain Diversification

Diffusion MRI tractography further suggests that long-range wiring organization represents a major axis of interspecific variation (22). In particular, the contrast between the relatively localized optic-lobe-associated trajectories observed in chick and the more broadly distributed trajectories observed in the gentoo penguin indicates that sensory routing architectures can differ substantially among species. The presence of a distinct lateral temporal cortex-like trajectory pattern in the penguin further supports the idea that ecological specialization may be accompanied by reorganization of pallial connectivity motifs.

At the same time, tractography-derived trajectory distributions and diffusion-based microstructural indices provided partially independent information, consistent with the broader observation that tractography and scalar diffusion metrics capture different aspects of tissue organization (22, 33, 34). For example, elevated cerebellar FA in the gentoo penguin was not accompanied by increased streamline representation within the same region, whereas dorsal pallium-associated trajectories were comparatively prominent. Inthe large-billed crow, by contrast, elevated FA was most evident in the cerebellum and optic lobe, whereas trajectory enrichment was concentrated in the anterior lobe. These comparisons underscore the complementary nature of tractography-derived trajectory distributions and diffusion-based microstructural measures and highlight the importance of interpreting both jointly in comparative analyses (22, 33).

### Histological Anchoring and Developmental Implications

A major strength of the present framework is that MRI-derived measurements were anchored to histological features. Nissl staining supported the anatomical boundaries identified by MRI, whereas myelin staining corresponded closely to major white-matter compartments and principal diffusion-derived pathways. This histological validation strengthens the biological interpretation of MRI-based volumetric and diffusion measurements and provides a foundation for identifying tissue properties that may contribute to interspecific variation in diffusion metrics, including myelination, axon caliber, axonal packing density, and orientation dispersion (22, 33). More broadly, linking MRI-derived phenotypes to underlying tissue properties creates new opportunities to connect comparative neuroanatomy with the developmental mechanisms that shape regional brain architecture (35, 36).

### A Case Study of Mosaic Specialization: The Penguin Brain

Penguins provide a particularly informative example of mosaic neuroanatomical specialization. In our dataset, the gentoo penguin exhibited a distinctive combination of relatively reduced AC prominence and pronounced cerebellar expansion, consistent with lineage-specific reallocation of neural investment. In birds, telencephalic interhemispheric connectivity relies on a limited number of commissural pathways, with the AC serving as the principal forebrain commissure. Tract-tracing studies have shown that AC projections originate from restricted arcopallial and amygdaloid sources and project broadly to contralateral sensorimotor and limbic targets, suggesting that variation in AC prominence may reflect differences in commissural routing rather than serving as a simple indicator of any particular sensory modality (17). These studies further indicate that, unlike in mammals, interhemispheric transfer of olfactory information in birds is not mediated primarily through the AC but may instead involve alternative diencephalic pathways such as the habenular commissure (17).

Accordingly, the reduced AC prominence observed in penguins should not be interpreted directly as evidence of olfactory specialization. Rather, it may reflect lineage-specific reorganization of commissural architecture associated with differences in the integration of bilateral sensory information across hemispheres. However, the present data do not allow us to distinguish among several non-mutually exclusive explanations, including altered reliance on non-AC commissural pathways, changes in interhemispheric sensory integration, or a greater emphasis on rapid unilateral processing.

The marked cerebellar expansion observed in penguins is likewise consistent with increased demands on visuovestibular integration, gaze stabilization, and precise sensorimotor timing. Although these characteristics are compatible with the requirements of three-dimensional underwater locomotion, they may not be unique to aquatic specialization and could instead reflect more general adaptations associated with high locomotor performance in birds. Together, these findings suggest that penguin brain evolution involved coordinated yet regionally selective reorganization of commissural and sensorimotor systems. The ecological and functional significance of these changes, however, remains to be evaluated through broader comparative analyses.

### Limitations and Future Directions

Several limitations should be considered when interpreting the present findings. Cross-species MRI comparisons remain influenced by variation in acquisition conditions, and diffusion tractography provides an indirect reconstruction of pathway organization rather than direct evidence of axonal connectivity (22). In addition, broader taxonomic sampling will be required to disentangle correlated ecological variables and determine whether the observed patterns reflect general principles of avian brain organization or lineage-specific specializations (18, 23).

For example, the penguin data suggest a distinctive combination of cerebellar organization, tectal projection patterns, and pallial trajectory architecture. However, additional sampling of both aquatic and nonaquatic species will be necessary to determine the extent to which these features are characteristic of aquatic adaptation more broadly. Despite these limitations, the present framework provides standardized measures of internal anatomical organization, quantitative volumetry, and whole-brain connectivity in a format suitable for comparative analyses. As such, it enables avian neuroanatomy to move beyond coarse comparisons of brain size toward explicit, testable hypotheses linking ecological demands, developmental programs, and internal circuit organization (36).

### Conclusion

We present a harmonized avian MRI resource together with a reproducible analytical framework for quantifying internal brain organization across diverse ecological contexts. By revealing mosaic variation in regional investment and long-range wiring architecture, while anchoring MRI-derived metrics to histological features, this study provides a scalable foundation for linking avian neuroanatomical diversity to ecological pressures, developmental programs, and evolutionary trajectories.

## Materials and Methods

### Brain Samples

We assembled a comparative MRI dataset comprising 16 avian species representing major avian clades and diverse ecological and behavioral categories, together with a flying mammalian outgroup, the Egyptian rousette bat, for selected diffusion-based comparisons. MRI datasets were obtained from two sources: (i) previously deposited datasets from the ABC and (ii) publicly available avian MRI datasets for canary, starling, pigeon, and zebra finch. Datasets were included when T2-weighted structural images provided sufficient contrast to delineate major brain compartments and, for diffusion analyses, when diffusion-weighted images possessed adequate angular and spatial resolution for tensor reconstruction and deterministic tractography. For publicly available datasets, original acquisition metadata were retained and integrated into a standardized metadata table.

### Image Preprocessing and Template Construction

Before analysis, T2-weighted images were inspected for orientation, field-of-view coverage, and major imaging artifacts. Non-brain tissue was removed manually or semi-automatically using 3D Slicer, after which brains were reoriented into a common anatomical convention based on corresponding coronal, sagittal, and horizontal planes. When required, image intensity was adjusted solely for visualization and identification of anatomical boundaries; all quantitative segmentations were performed using the original structural image volumes. Species-specific templates were generated from the available structural images for each species and used as reference spaces for regional parcellation. For cross-species visualization, segmented regions were exported as 3D surface models and rendered from standardized viewpoints. Manual reorientation was performed in 3D Slicer (version 5.8.1) using the Transforms module, with the anterior commissure and midsagittal plane used as anatomical landmarks. Only rigid linear transformations, consisting of translations and rotations, were applied.

### Anatomical Segmentation and Volumetric Analysis

T2-weighted images were imported into 3D Slicer, and anatomical segmentation was performed using the Segment Editor module. First, multiple consecutive slices were coarsely labeled with the Paint tool to distinguish the whole brain (“brain”) from surrounding non-brain tissue (“other”). The whole brain was then extracted as a single segment using the Grow from Seeds algorithm. For regional parcellation, anatomical boundaries were delineated and filled using the Draw tool on every other slice, and intermediate slices were generated by interpolation with Fill between Slices to produce 3D segmentations. Segment volumes were calculated using the Segment Statistics module. Volumes (mm³) were derived from the number of voxels contained within each segment, and the proportional volume of each region was expressed relative to total brain volume.

### Anterior Commissure (AC) Measurement

The AC was identified on coronal T2-weighted sections as a transverse commissural fiber bundle located ventral to the telencephalic midline within the rostral forebrain/preoptic region. For each species, AC area was measured on the coronal section exhibiting the largest cross-sectional profile of the commissure. The AC was manually outlined as a region of interest, and the corresponding whole-brain cross-sectional area was measured on the same section. Relative AC size was calculated as the ratio of AC area to whole-brain area. Allometric relationships were evaluated by comparing AC area with whole-brain area across species. For each species, AC area and whole-brain cross-sectional area were measured once on a single coronal section corresponding to the maximal cross-sectional profile of the AC.

### Cerebellar Foliation Analysis

Cerebellar foliation was assessed using midsagittal or near-midsagittal T2-weighted sections and, when available, corresponding histological sections. The primary cerebellar arbor was traced from the central white-matter core, and folial branches were categorized as either basal or terminal branches. Basal branches were defined as primary branches arising from the central cerebellar white-matter trunk, whereas terminal branches were defined as distal folial endings visible at the pial surface. Branch numbers were quantified from equivalent anatomical planes across species whenever possible. Among the 14 bird species examined, cerebellar foliation was analyzed primarily using midsagittal T2-weighted images. The Humboldt penguin was excluded because the cerebellar folial architecture could not be reliably identified in the available MRI data. For the domestic chicken, branch counts were performed using an existing chicken brain atlas, as suitable midsagittal MRI sections were not available. All counts were performed by a single observer.

### Histological Validation

To validate MRI-derived anatomical boundaries, histological analyses were performed in three representative species: the large-billed crow, gentoo penguin, and mandarin duck. Brains were sectioned in planes corresponding to the MRI images and stained using Nissl staining to visualize cytoarchitecture and Luxol fast blue (LFB) staining to identify myelinated fiber-rich compartments.

Histological sections were prepared at a thickness of 50–100 µm. Before Nissl and LFB staining, sections were defatted. Briefly, sections were immersed twice in distilled water for 2 min each and then in 70% ethanol for 2 min. Sections were subsequently immersed in 100% ethanol five times for 2 min each, followed by xylene for 5 min. After defatting, sections were rehydrated through four changes of 100% ethanol for 2 min each and then processed for either LFB or Nissl staining.

For LFB staining, defatted slides were immersed in 95% ethanol for 1 min. Luxol fast blue was dissolved in 95% ethanol at a final concentration of 0.2%, and the slides were incubated in this solution at 60°C for 16–24 h. After the staining solution had cooled to room temperature, the slides were removed and immersed in 95% ethanol for 2 min. The slides were then rinsed under running tap water and immersed in distilled water for 5 min. Differentiation was performed by gently agitating the slides in 0.05% lithium carbonate for 5–15 s, followed by agitation in 70% ethanol for 2 min. The slides were further differentiated in fresh 70% ethanol twice for 1 min each. After differentiation, the slides were washed under running tap water, immersed in distilled water for 5 min, dehydrated, cleared, and mounted.

For Nissl staining, defatted slides were immersed in 70% ethanol for 2 min, rinsed under running tap water, and immersed in distilled water for 5 min. The slides were then stained in 0.5% cresyl violet acetate for 6 min. Differentiation was performed by dipping the slides twice in 70% ethanol containing 0.1% acetic acid, followed by three dips in 95% ethanol containing 0.1% acetic acid. The solution was then replaced, and the slides were dipped three additional times in fresh 95% ethanol containing 0.1% acetic acid. The slides were subsequently dipped 10 times in 100% ethanol and transferred to fresh 100% ethanol for 2 min before dehydration, clearing, and mounting.

For dehydration and clearing, stained slides were immersed in 100% ethanol for 2 min and then cleared in xylene three times for 5 min each. Sections were mounted using a xylene-based mounting medium. Sections used for analysis were selected based on anatomical landmarks of the cerebrum and midbrain. Images were acquired using a Keyence BZ-X800 microscope.

MRI and histological sections were compared at anterior, middle, and posterior levels. Anatomical boundaries were evaluated based on the correspondence between MRI contrast, cytoarchitectonic transitions, and myelin-rich compartments.

### Diffusion/Tractography Analysis

Diffusion-weighted images were reconstructed using the DTI model implemented in DSI Studio (https://dsi-studio.labsolver.org/). Diffusion tensors were estimated for each voxel, and scalar diffusion metrics, including FA, were derived from the resulting tensor model. Tractography was performed using the generalized deterministic tracking algorithm implemented in DSI Studio. For whole-brain tractography, 10,000,000 streamlines were generated using an FA threshold of 0.1, a minimum streamline length of 1 mm, and a maximum streamline length of 300 mm. Random maximum angle and trilinear interpolation settings were applied as global tracking parameters. The resulting streamlines were subsequently classified into ten clusters using the K-means clustering algorithm as implemented in DSI Studio.

### Statistical Analysis

Morphometric and diffusion-derived measurements were analyzed as exploratory cross-species comparisons. AC cross-sectional area was compared with whole-brain cross-sectional area to evaluate broad allometric scaling across species, and relative AC size was calculated as the ratio of AC area to whole-brain cross-sectional area. Cerebellar foliation was quantified by counting basal and terminal branches from comparable midsagittal or near-midsagittal sections. Segmentation-derived regional volumes were normalized to total brain volume and expressed as proportional volume fractions. Diffusion-derived metrics, including streamline distributions and FA values, were compared across representative species to identify conserved and divergent anatomical patterns. Because the sample size and data sources varied across species, these analyses were treated as descriptive and hypothesis-generating rather than formal phylogenetically corrected tests.

## Acknowledgments

The authors would like to thank Drs. Iguchi and Seki, who prepared Nissl and LFB staining samples (TMiMs). This work was supported by a Leading Initiative for Excellent Young Researchers (LEADER) grant (grant number 2020L0019), JSPS KAKENHI grants (20K22665, 22H02638, 23K23901), the FY2021 Research Grant from the Takeda Science Foundation (to T.K.). C.O-M. was supported by JSPS KAKENHI-Grants (25K02289), AMED under Grant Number JP24gm1310012. T.T. was supported by JSPS KAKENHI-Grants (25K02565 and 21K19464). ChatGPT (OpenAI, GPT-5.5 Thinking) was used to assist with English language editing during manuscript preparation.

## Supplementary Figures

**Fig. S1.**
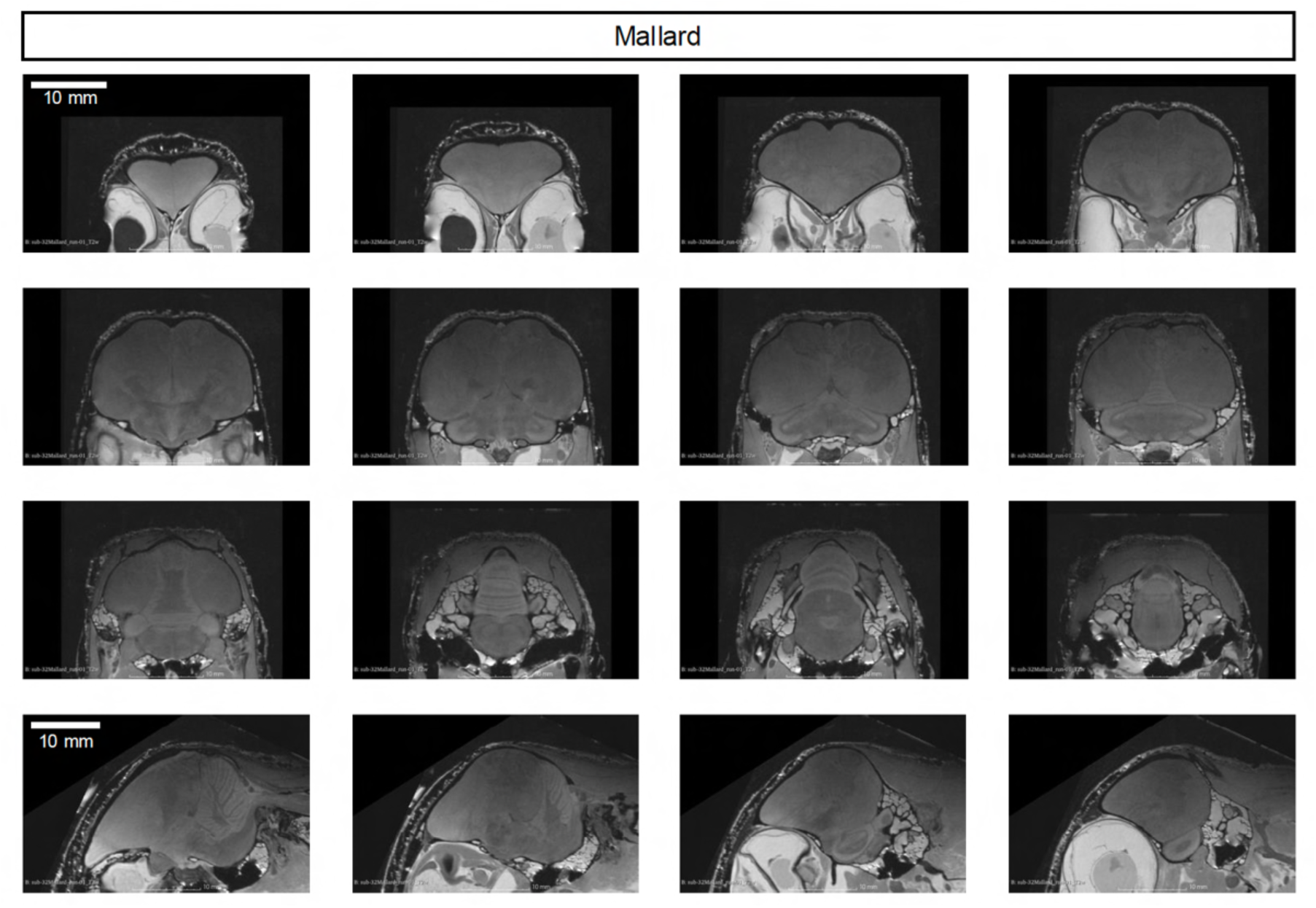
Serial coronal and sagittal T2-weighted MRI sections of the mallard. Representative serial T2-weighted MRI sections of the mallard brain are shown to provide an overview of whole-brain morphology. Consecutive coronal sections are shown together with sagittal sections to illustrate the anatomical continuity of the forebrain, midbrain, cerebellum, and brainstem. Scale bars, 10 mm.

**Fig. S2.**
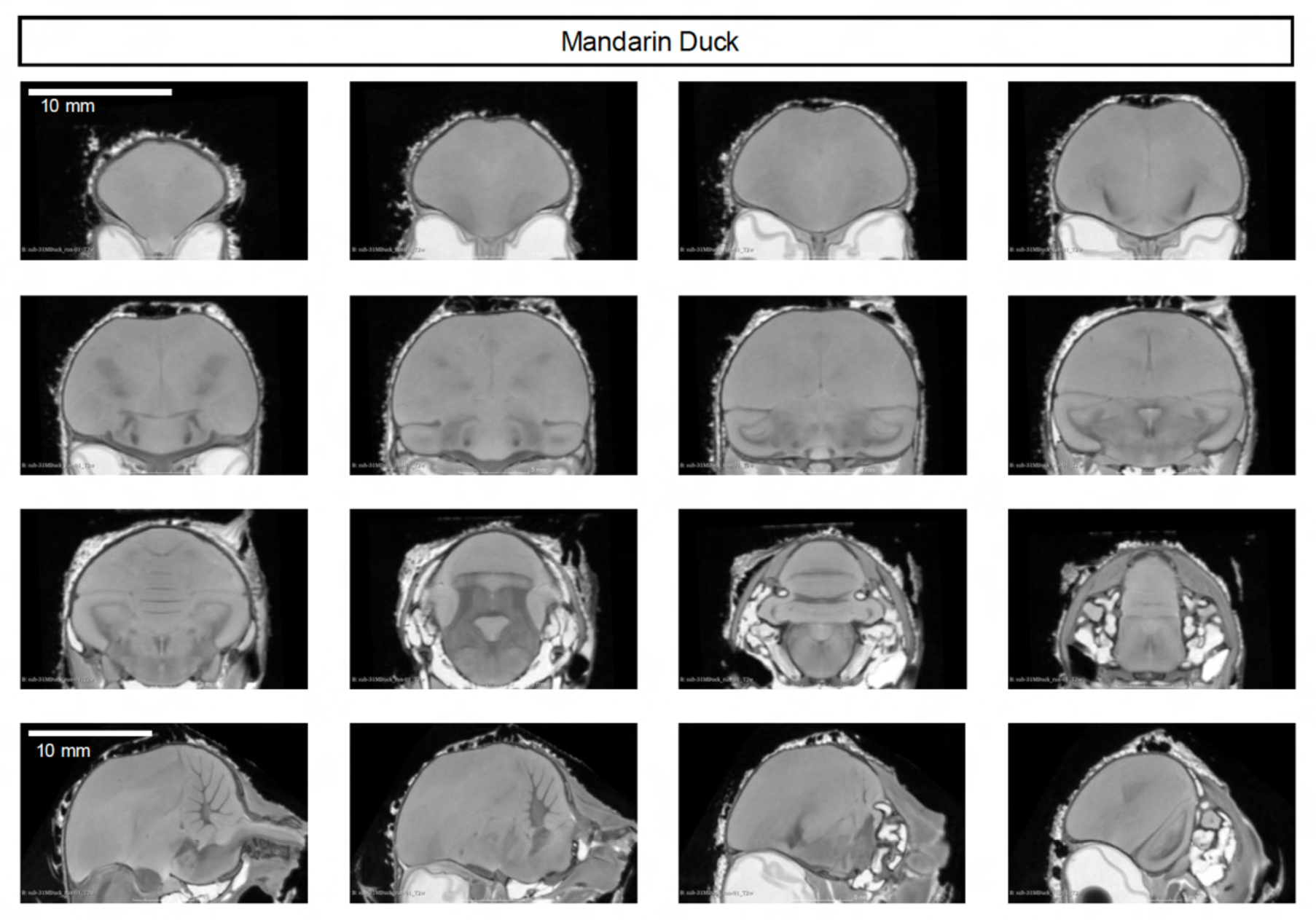
Serial coronal and sagittal T2-weighted MRI sections of the mandarin duck. Representative serial T2-weighted MRI sections of the mandarin duck brain are shown to provide an overview of whole-brain morphology. Consecutive coronal sections are shown together with sagittal sections to illustrate the anatomical continuity of the forebrain, midbrain, cerebellum, and brainstem. Scale bars, 10 mm.

**Fig. S3.**
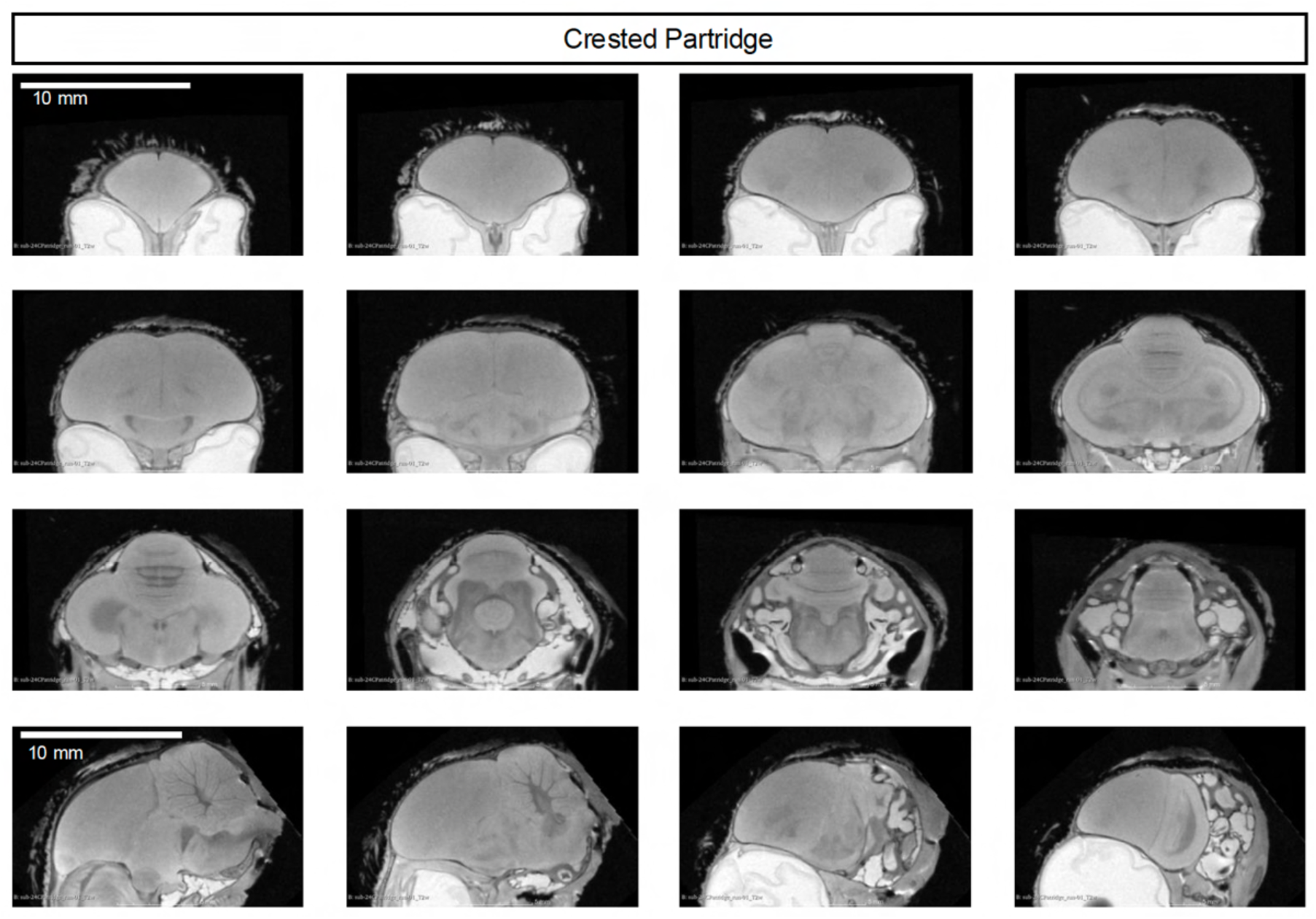
Serial coronal and sagittal T2-weighted MRI sections of the crested partridge. Representative serial T2-weighted MRI sections of the crested partridge brain are shown to provide an overview of whole-brain morphology. Consecutive coronal sections are shown together with sagittal sections to illustrate the anatomical continuity of the forebrain, midbrain, cerebellum, and brainstem. Scale bars, 10 mm.

**Fig. S4.**
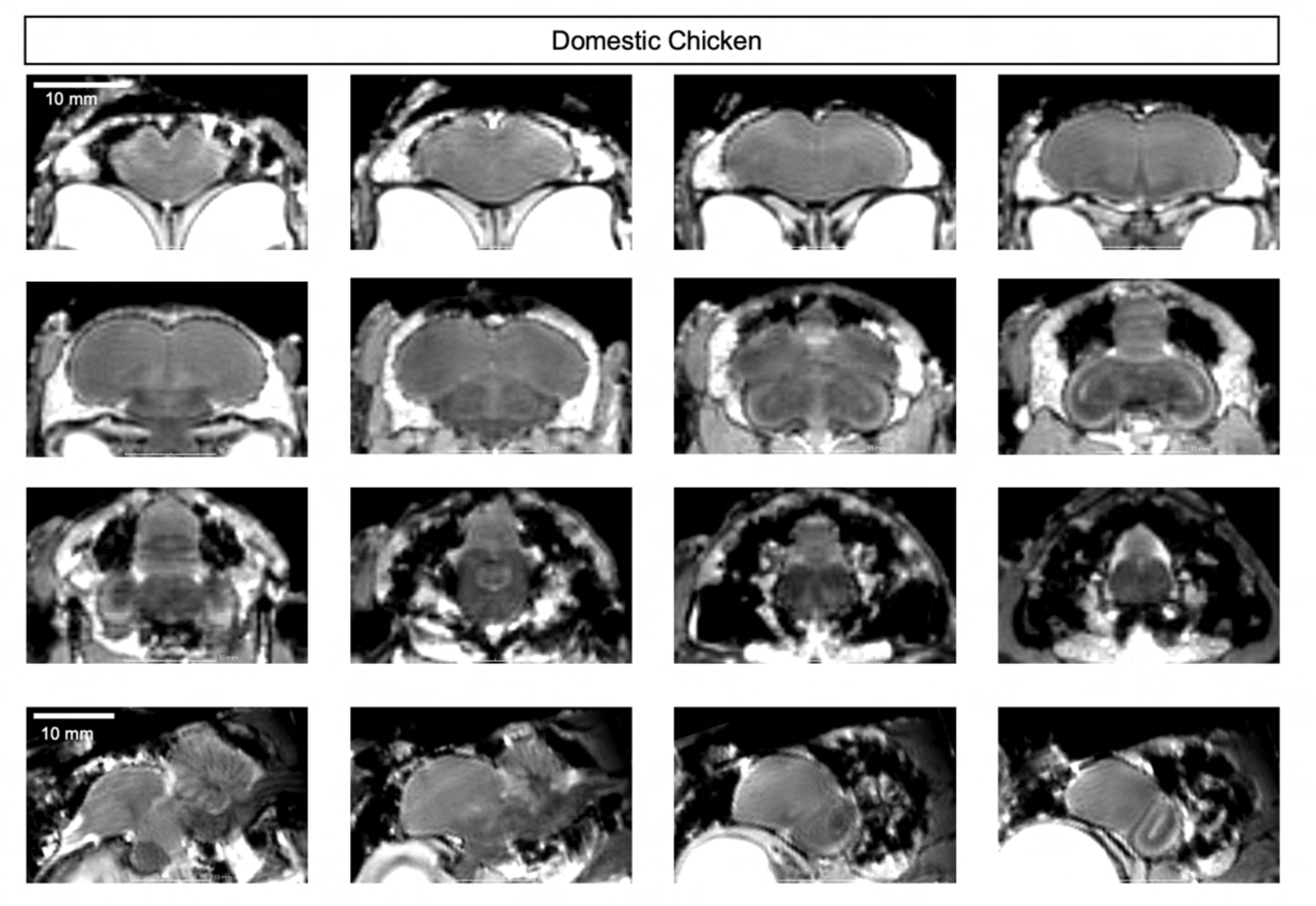
Serial coronal and sagittal T2-weighted MRI sections of the domestic chicken. Representative serial T2-weighted MRI sections of the domestic chicken brain are shown to provide an overview of whole-brain morphology. Consecutive coronal sections are shown together with sagittal sections to illustrate the anatomical continuity of the forebrain, midbrain, cerebellum, and brainstem. Scale bars, 10 mm.

**Fig. S5.**
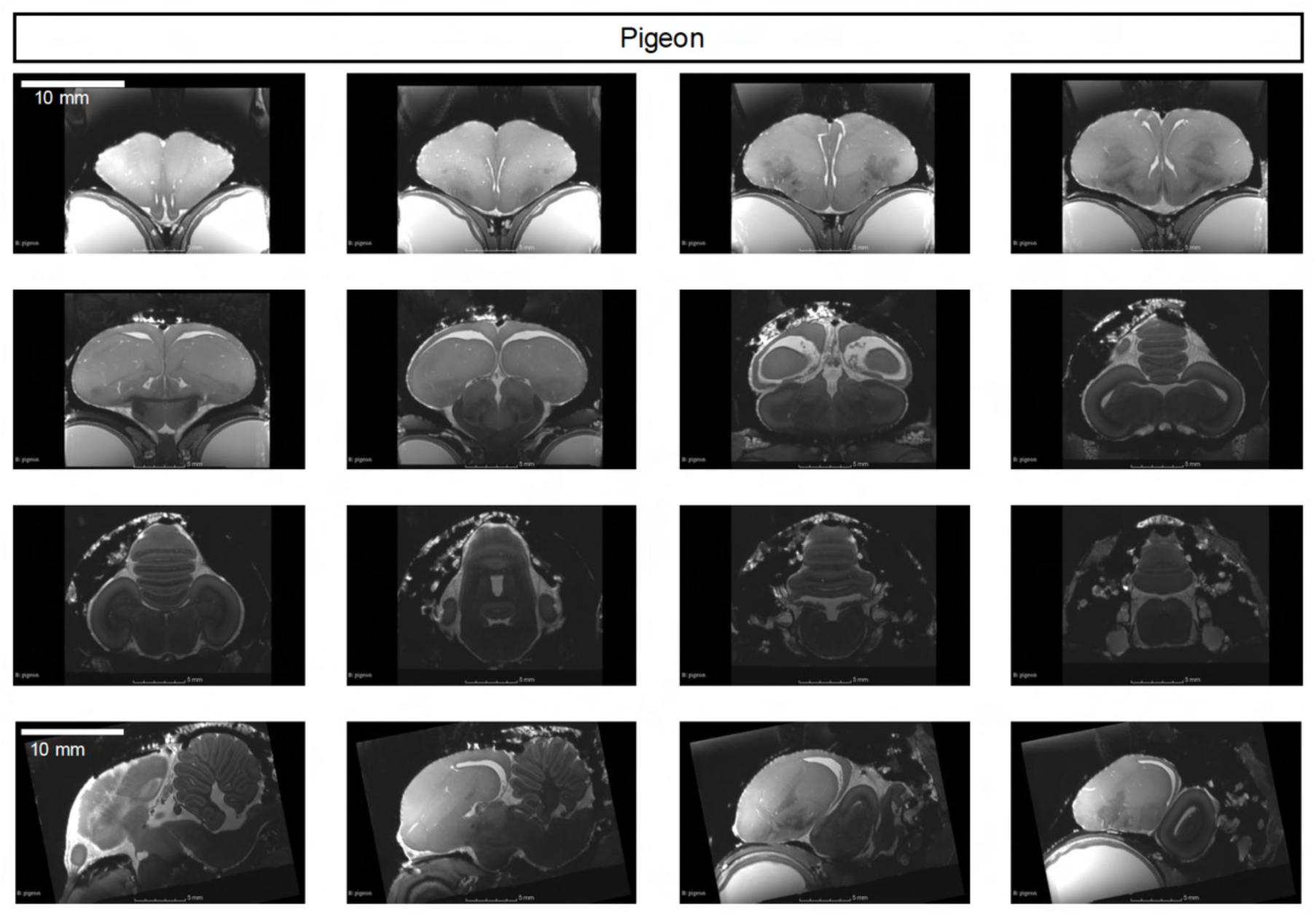
Serial coronal and sagittal T2-weighted MRI sections of the pigeon. Representative serial T2-weighted MRI sections of the pigeon brain are shown to provide an overview of whole-brain morphology. Consecutive coronal sections are shown together with sagittal sections to illustrate the anatomical continuity of the forebrain, midbrain, cerebellum, and brainstem. Scale bars, 10 mm.

**Fig. S6.**
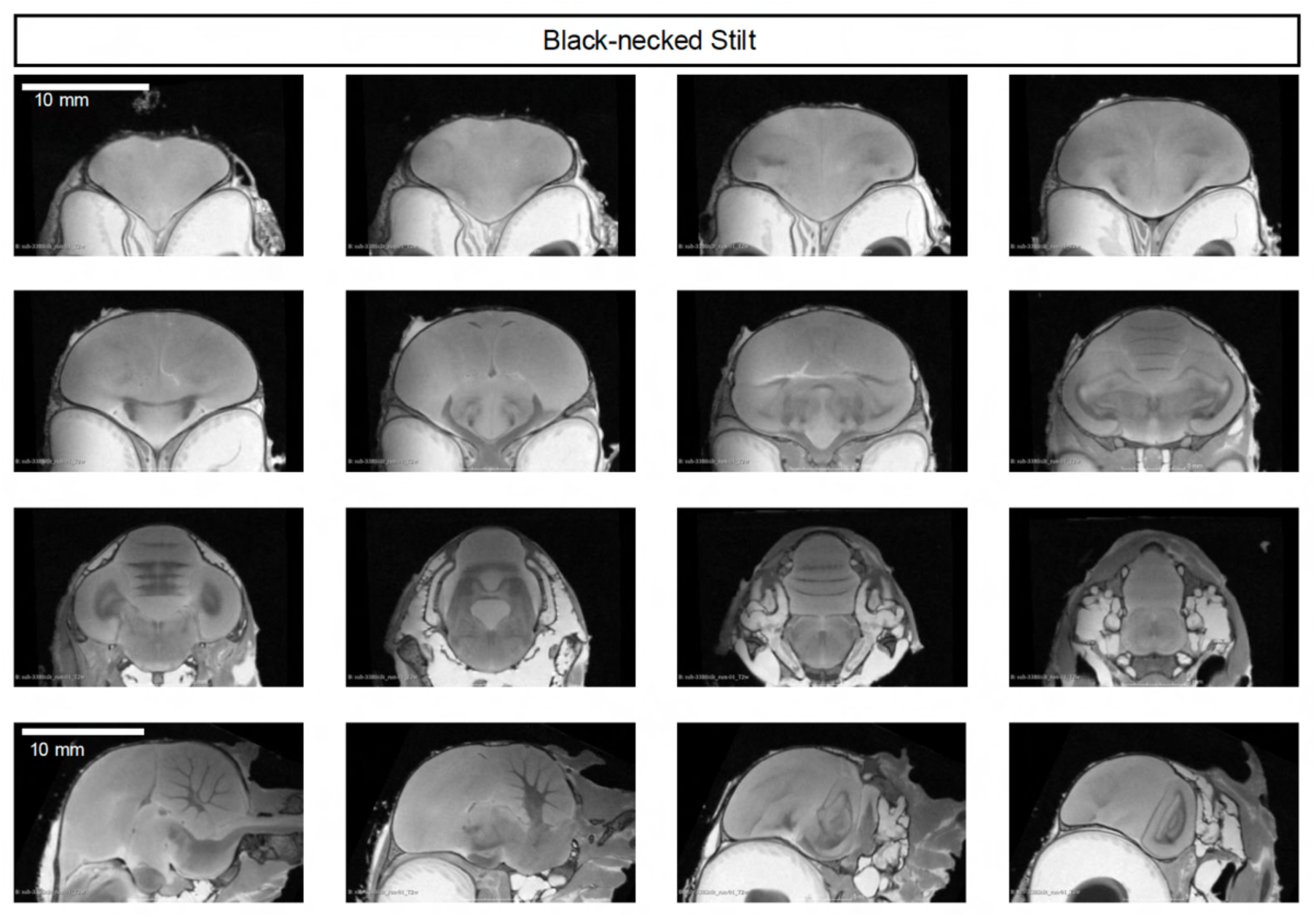
Serial coronal and sagittal T2-weighted MRI sections of the black-necked stilt. Representative serial T2-weighted MRI sections of the black-necked stilt brain are shown to provide an overview of whole-brain morphology. Consecutive coronal sections are shown together with sagittal sections to illustrate the anatomical continuity of the forebrain, midbrain, cerebellum, and brainstem. Scale bars, 10 mm.

**Fig. S7.**
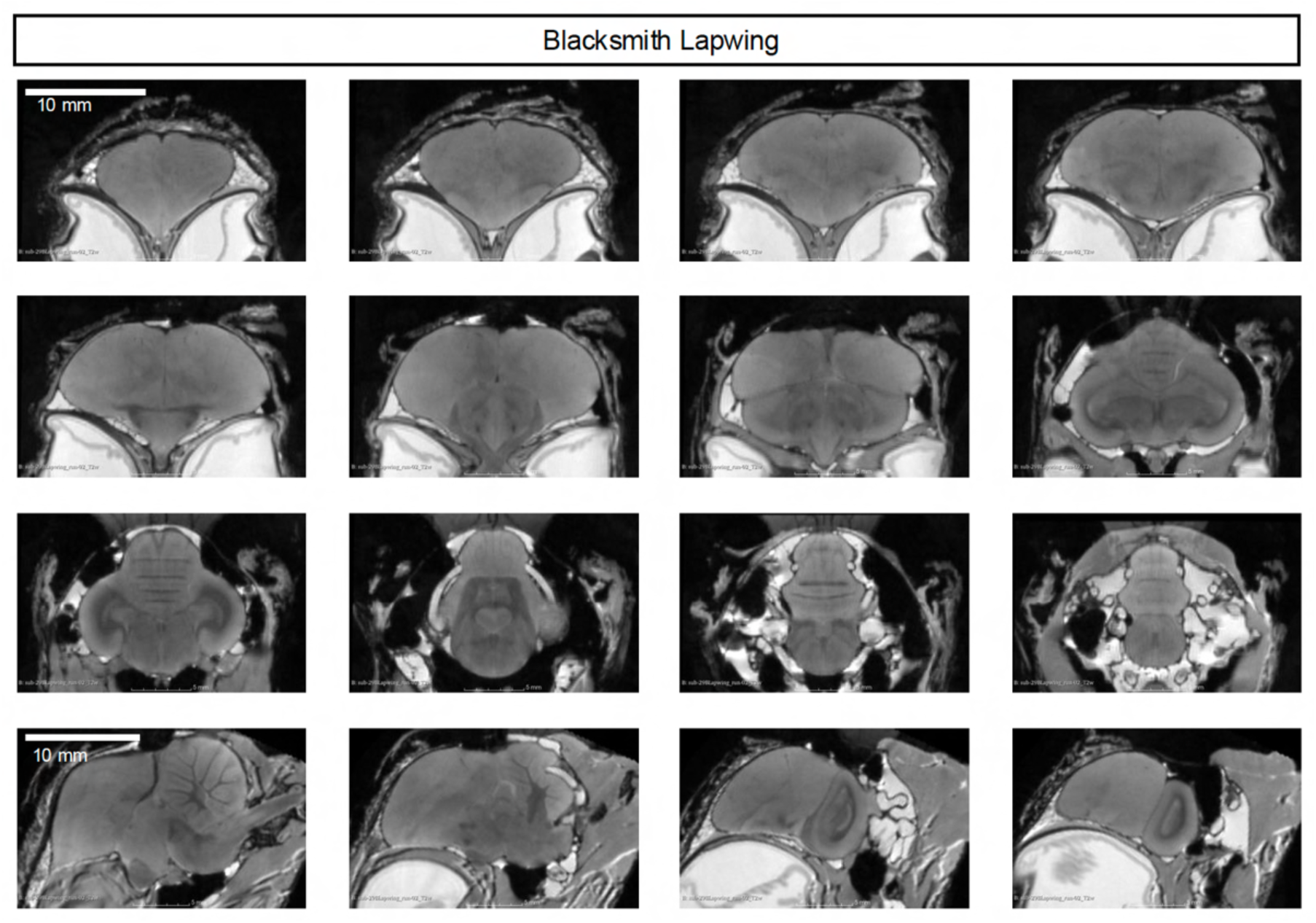
Serial coronal and sagittal T2-weighted MRI sections of the blacksmith lapwing. Representative serial T2-weighted MRI sections of the blacksmith lapwing brain are shown to provide an overview of whole-brain morphology. Consecutive coronal sections are shown together with sagittal sections to illustrate the anatomical continuity of the forebrain, midbrain, cerebellum, and brainstem. Scale bars, 10 mm.

**Fig. S8.**
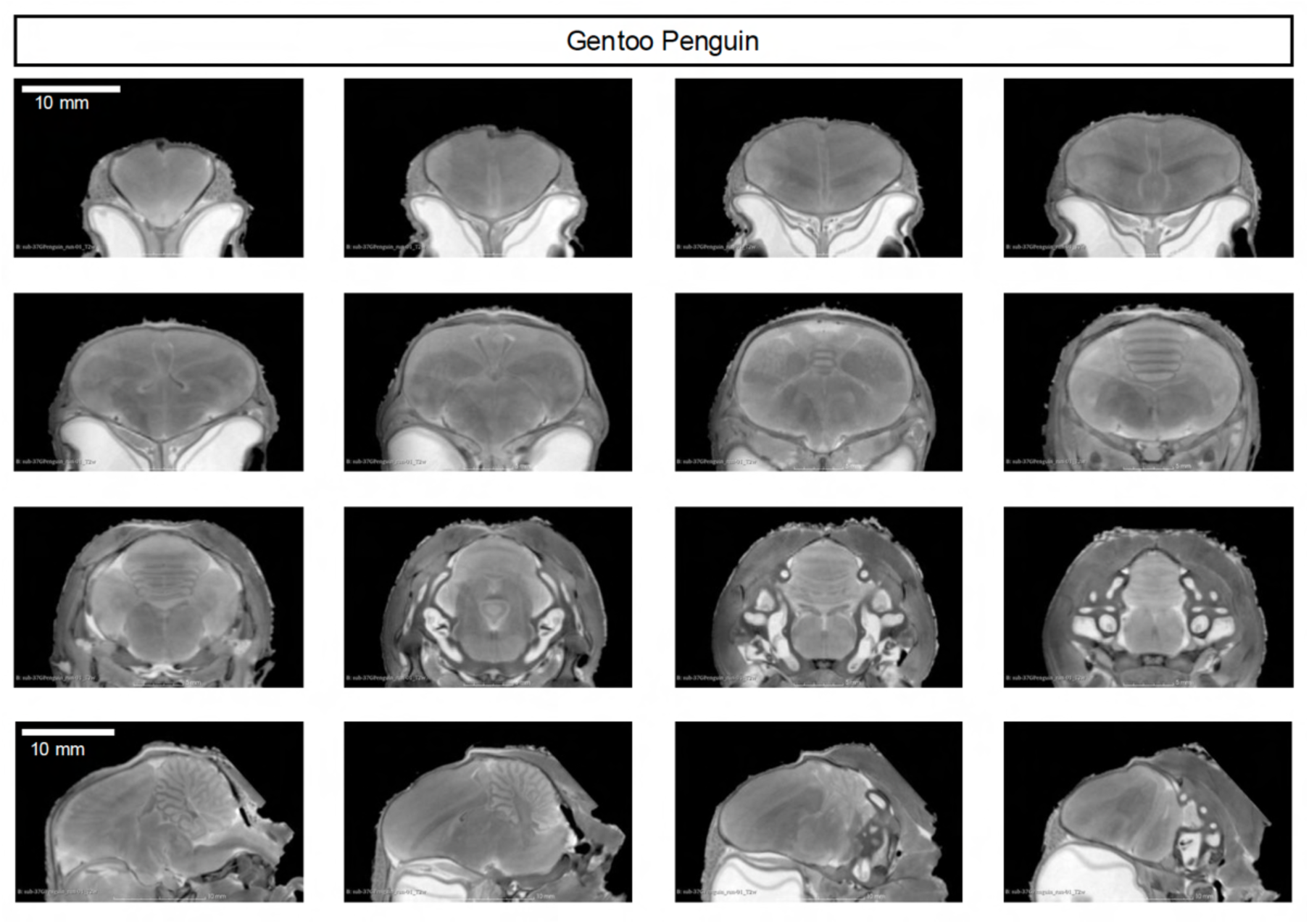
Serial coronal and sagittal T2-weighted MRI sections of the gentoo penguin. Representative serial T2-weighted MRI sections of the gentoo penguin brain are shown to provide an overview of whole-brain morphology. Consecutive coronal sections are shown together with sagittal sections to illustrate the anatomical continuity of the forebrain, midbrain, cerebellum, and brainstem. Scale bars, 10 mm.

**Fig. S9.**
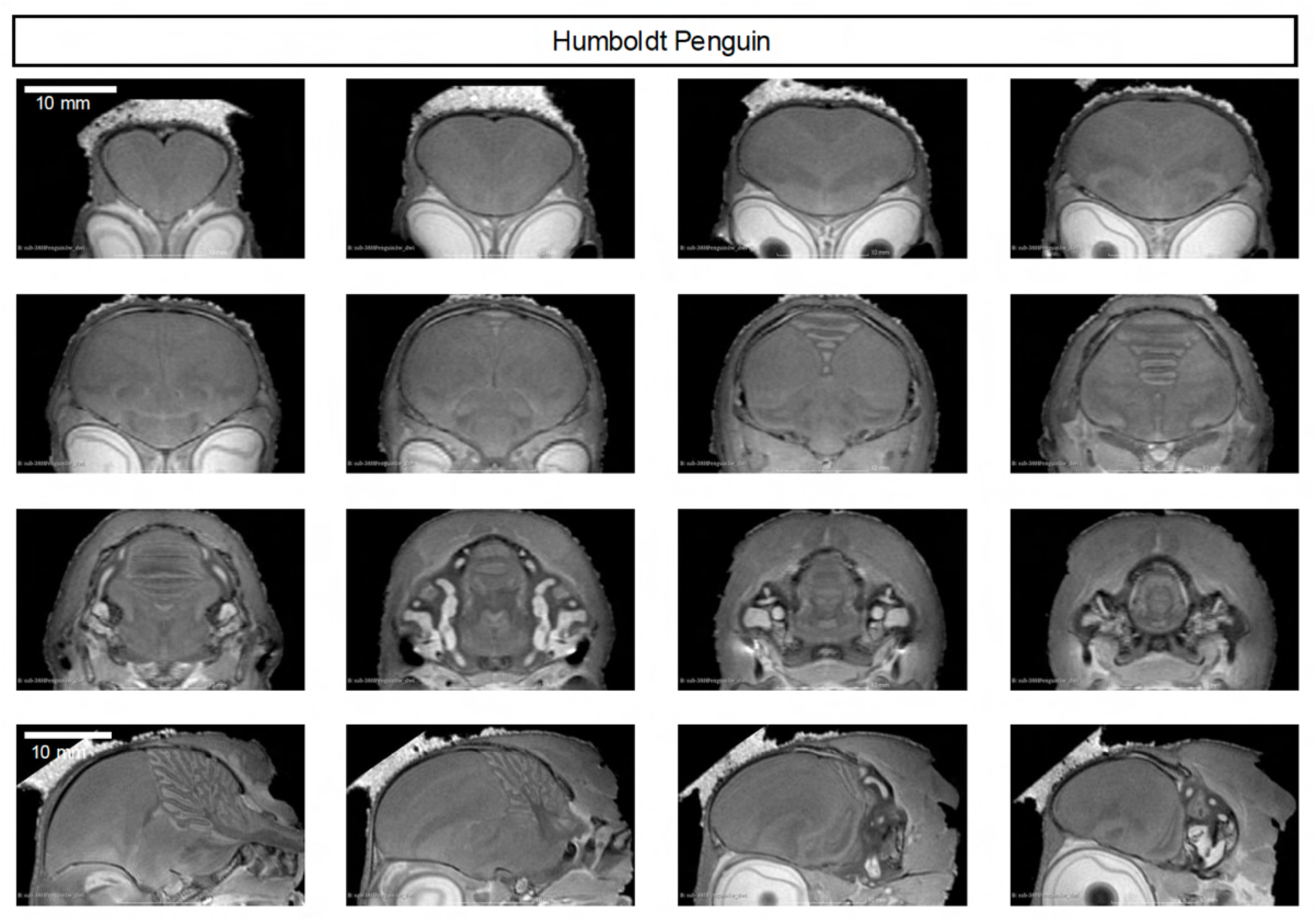
Serial coronal and sagittal T2-weighted MRI sections of the Humboldt penguin. Representative serial T2-weighted MRI sections of the Humboldt penguin brain are shown to provide an overview of whole-brain morphology. Consecutive coronal sections are shown together with sagittal sections to illustrate the anatomical continuity of the forebrain, midbrain, cerebellum, and brainstem. Scale bars, 10 mm.

**Fig. S10.**
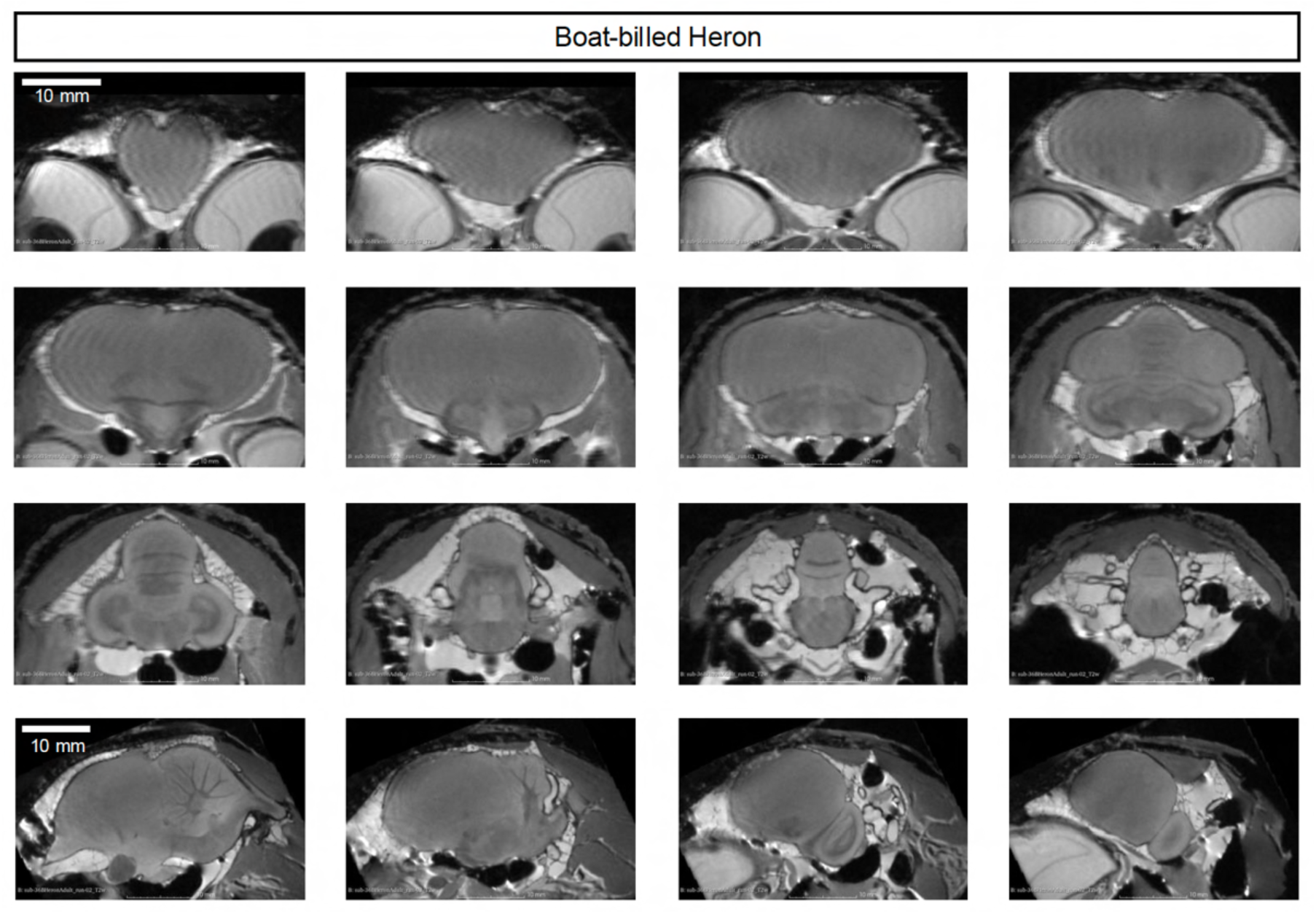
Serial coronal and sagittal T2-weighted MRI sections of the boat-billed heron. Representative serial T2-weighted MRI sections of the boat-billed heron brain are shown to provide an overview of whole-brain morphology. Consecutive coronal sections are shown together with sagittal sections to illustrate the anatomical continuity of the forebrain, midbrain, cerebellum, and brainstem. Scale bars, 10 mm.

**Fig. S11.**
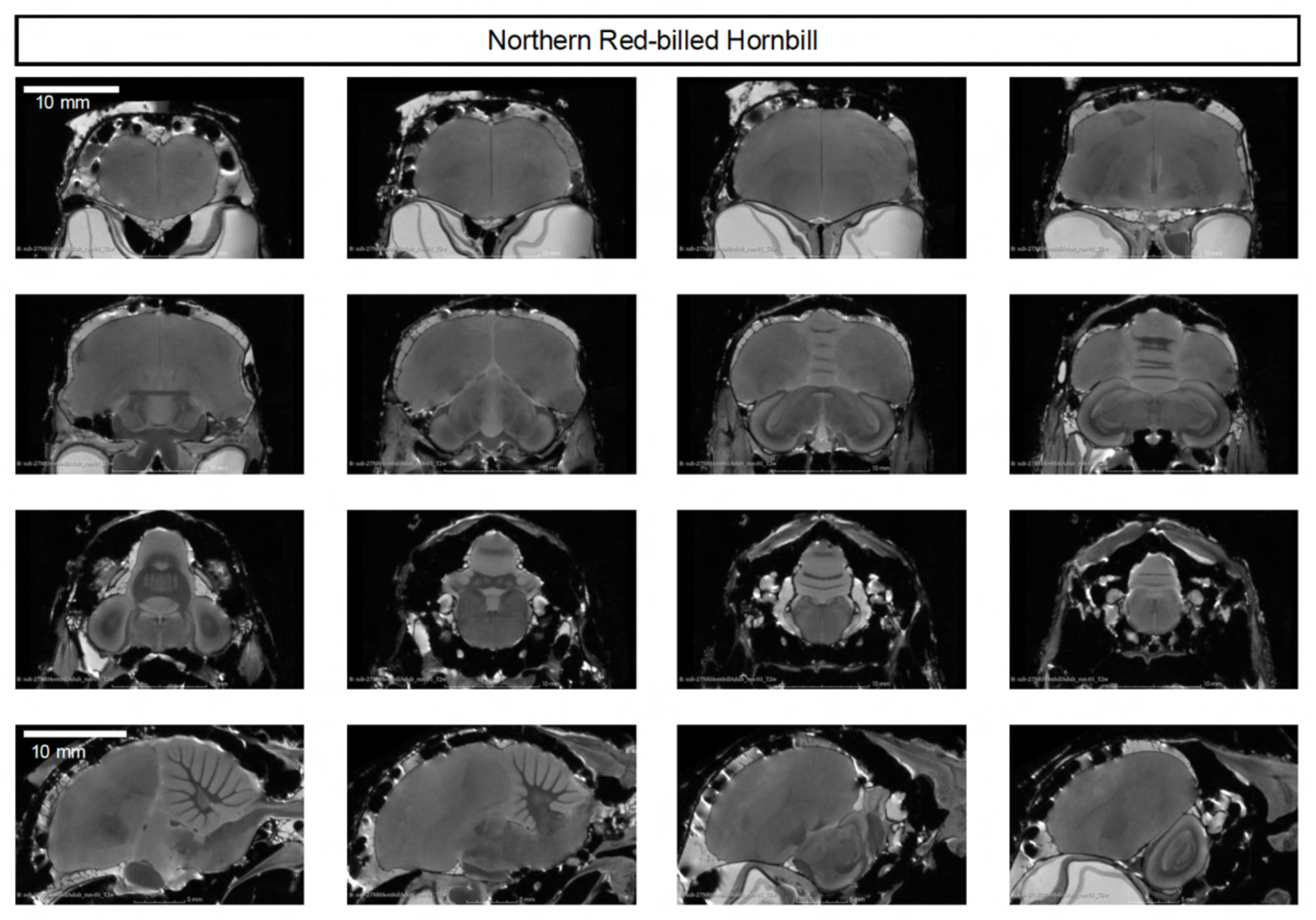
Serial coronal and sagittal T2-weighted MRI sections of the northern red-billed hornbill. Representative serial T2-weighted MRI sections of the northern red-billed hornbill brain are shown to provide an overview of whole-brain morphology. Consecutive coronal sections are shown together with sagittal sections to illustrate the anatomical continuity of the forebrain, midbrain, cerebellum, and brainstem. Scale bars, 10 mm.

**Fig. S12.**
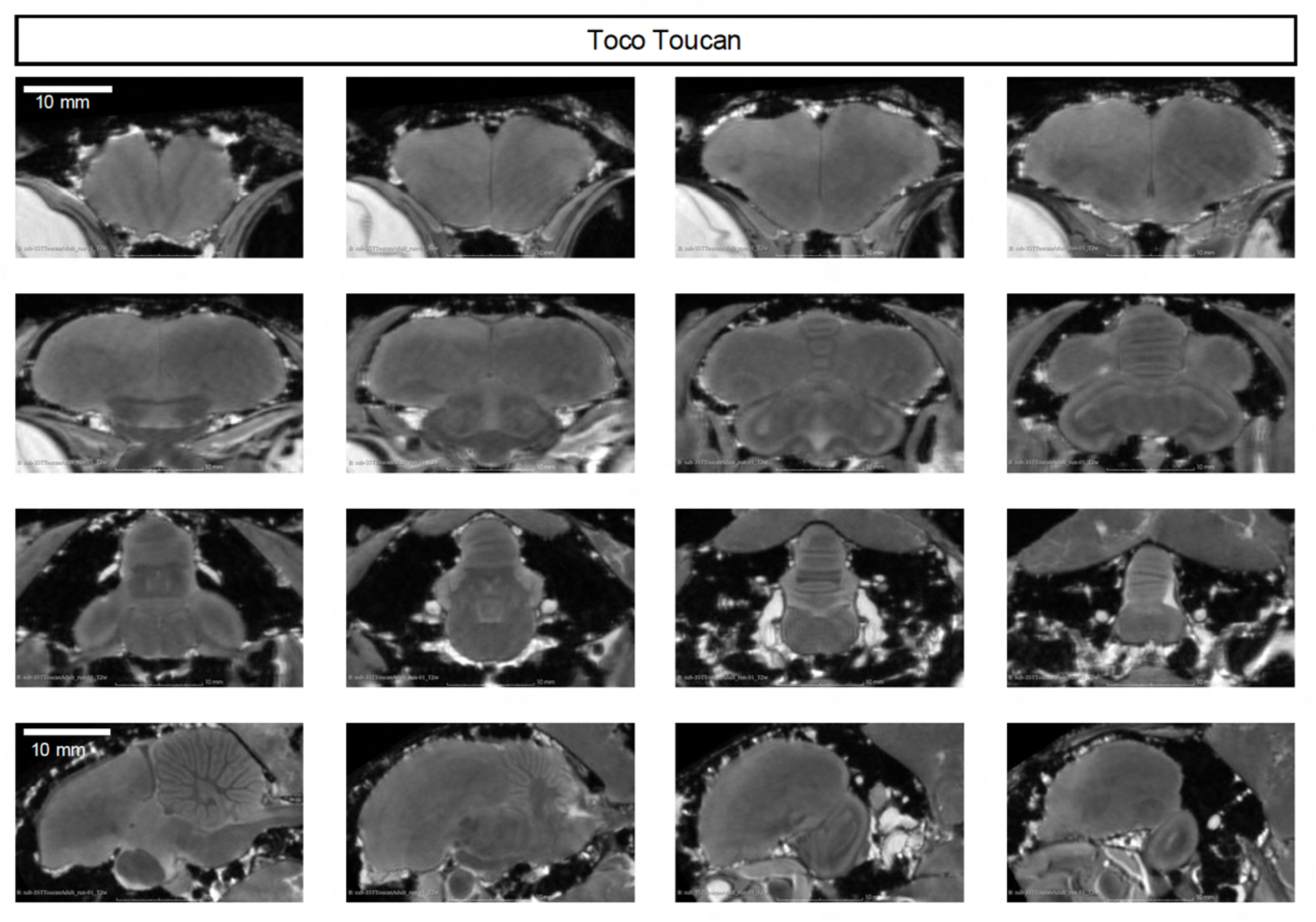
Serial coronal and sagittal T2-weighted MRI sections of the toco toucan. Representative serial T2-weighted MRI sections of the toco toucan brain are shown to provide an overview of whole-brain morphology. Consecutive coronal sections are shown together with sagittal sections to illustrate the anatomical continuity of the forebrain, midbrain, cerebellum, and brainstem. Scale bars, 10 mm.

**Fig. S13.**
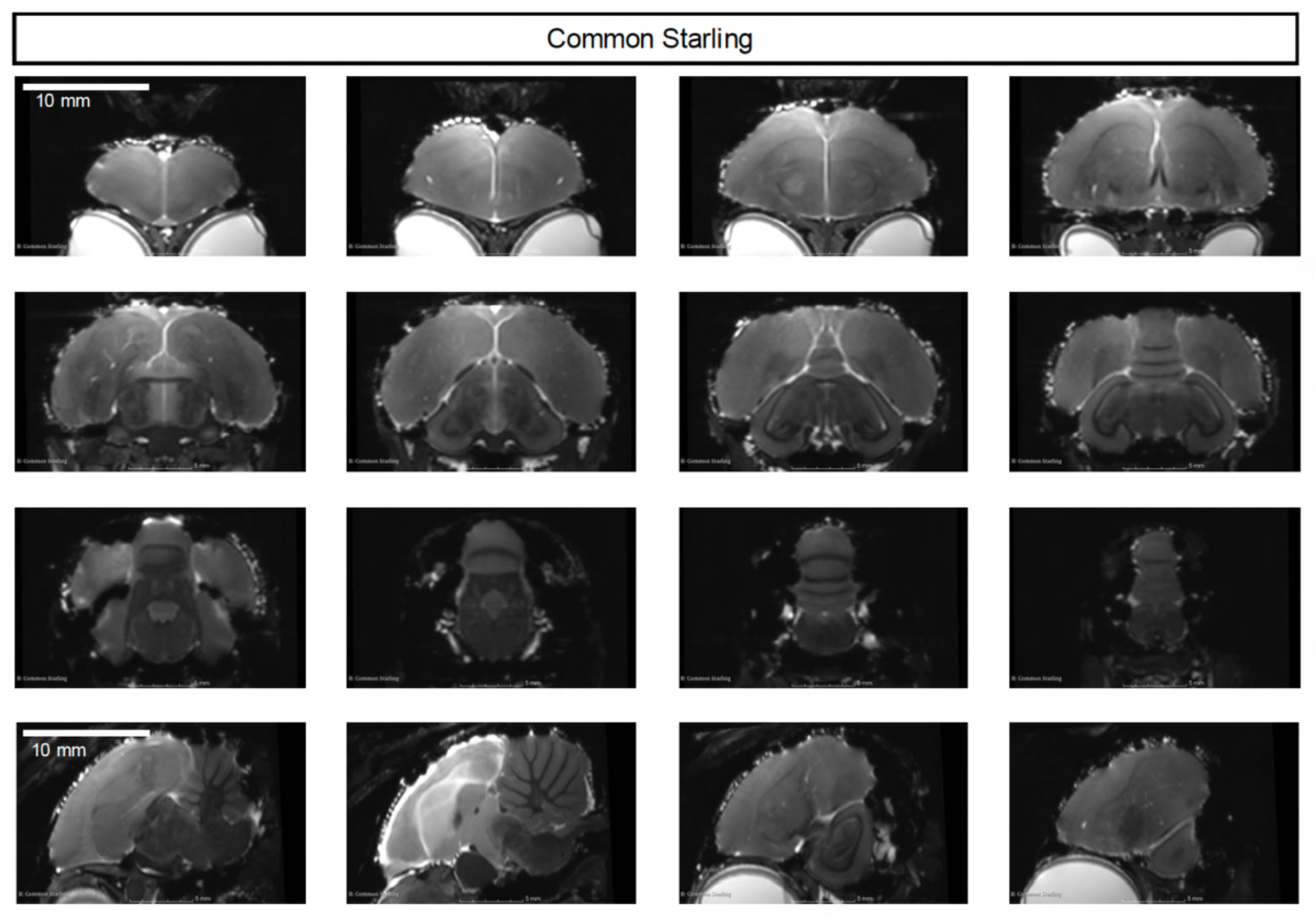
Serial coronal and sagittal T2-weighted MRI sections of the common starling. Representative serial T2-weighted MRI sections of the common starling brain are shown to provide an overview of whole-brain morphology. Consecutive coronal sections are shown together with sagittal sections to illustrate the anatomical continuity of the forebrain, midbrain, cerebellum, and brainstem. Scale bars, 10 mm.

**Fig. S14.**
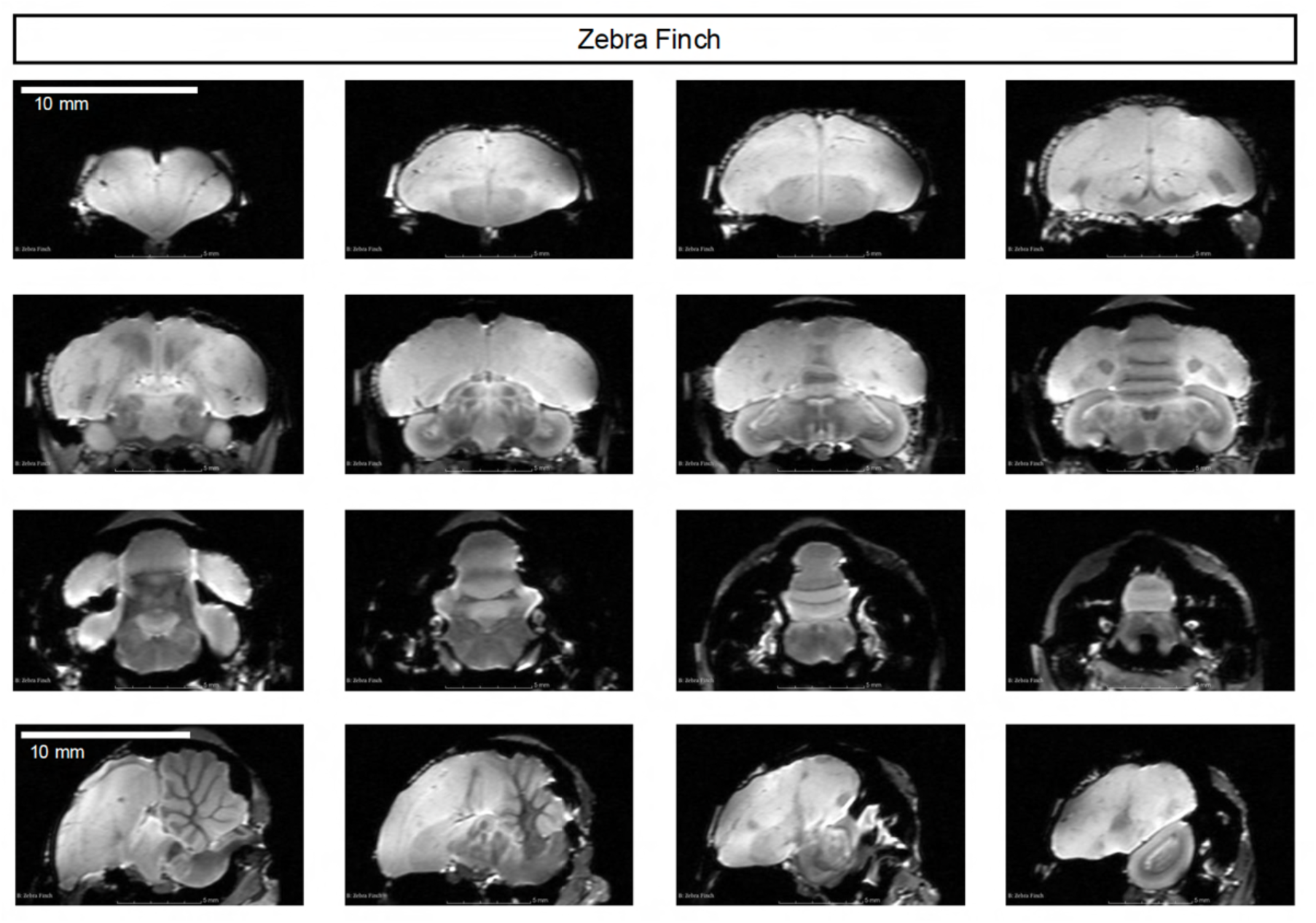
Serial coronal and sagittal T2-weighted MRI sections of the zebra finch. Representative serial T2-weighted MRI sections of the zebra finch brain are shown to provide an overview of whole-brain morphology. Consecutive coronal sections are shown together with sagittal sections to illustrate the anatomical continuity of the forebrain, midbrain, cerebellum, and brainstem. Scale bars, 10 mm.

**Fig. S15.**
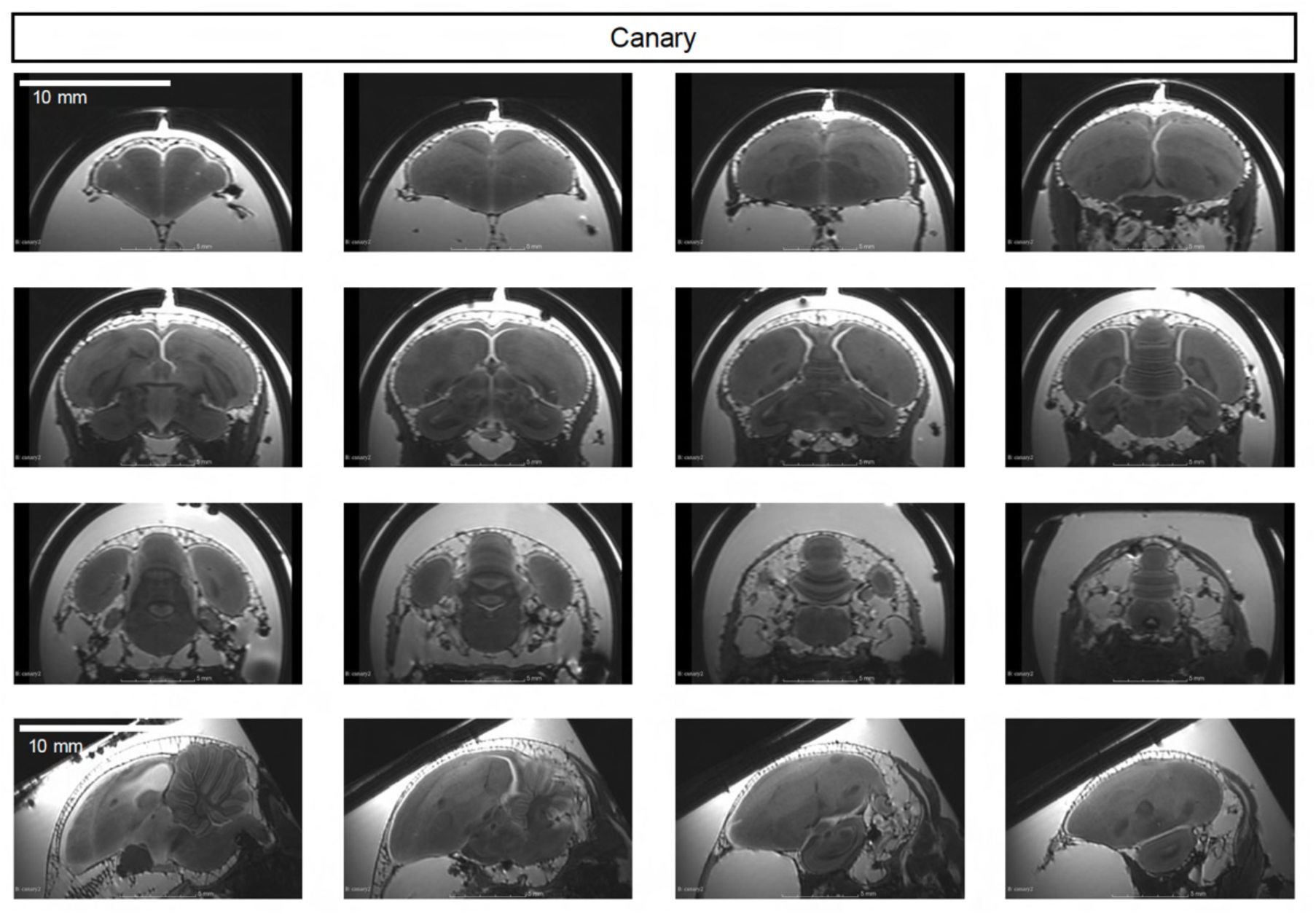
Serial coronal and sagittal T2-weighted MRI sections of the canary. Representative serial T2-weighted MRI sections of the canary brain are shown to provide an overview of whole-brain morphology. Consecutive coronal sections are shown together with sagittal sections to illustrate the anatomical continuity of the forebrain, midbrain, cerebellum, and brainstem. Scale bars, 10 mm.

**Fig. S16.**
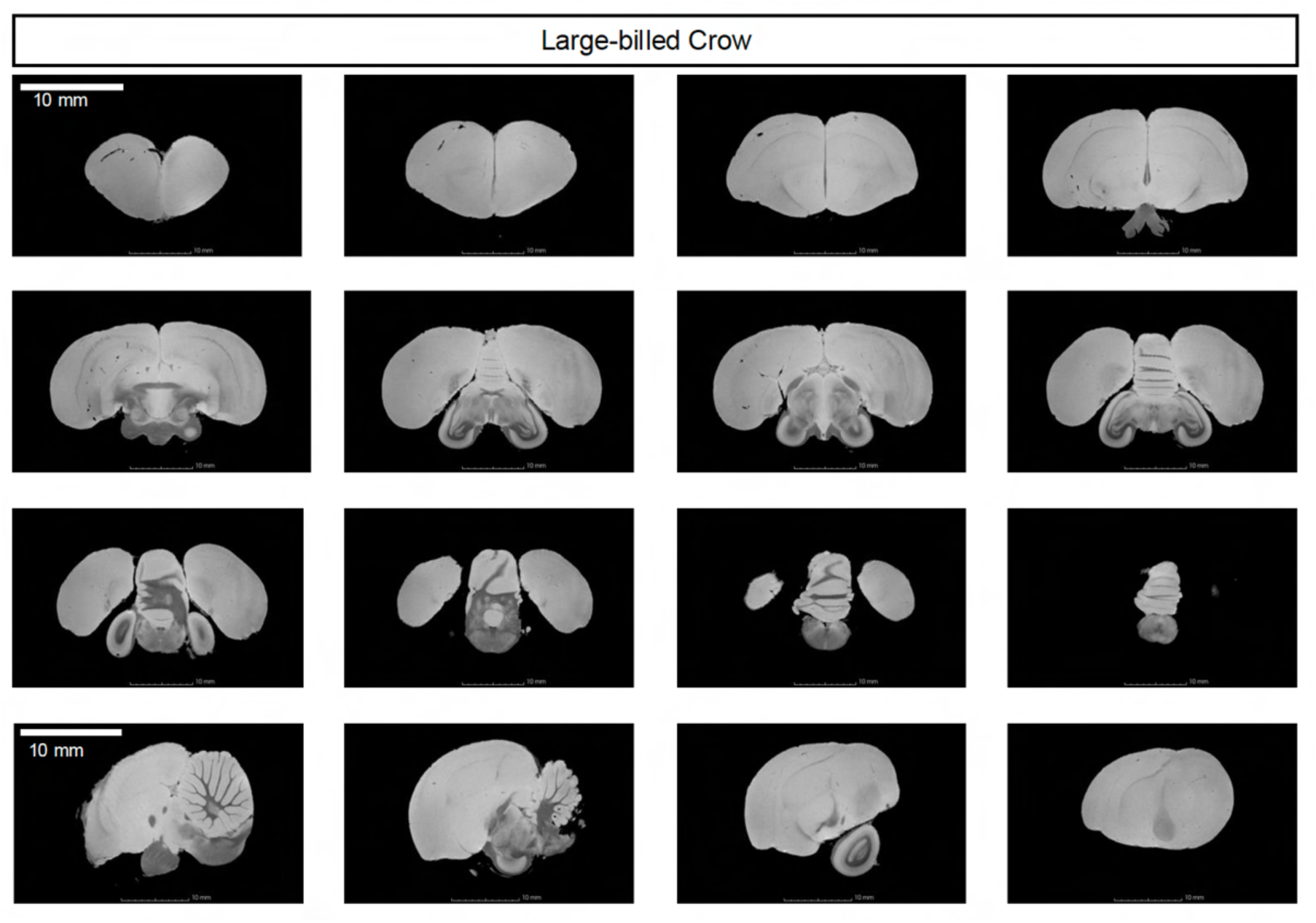
Serial coronal and sagittal T2-weighted MRI sections of the large-billed crow. Representative serial T2-weighted MRI sections of the large-billed crow brain are shown to provide an overview of whole-brain morphology. Consecutive coronal sections are shown together with sagittal sections to illustrate the anatomical continuity of the forebrain, midbrain, cerebellum, and brainstem. Scale bars, 10 mm.

**Fig. S17.**
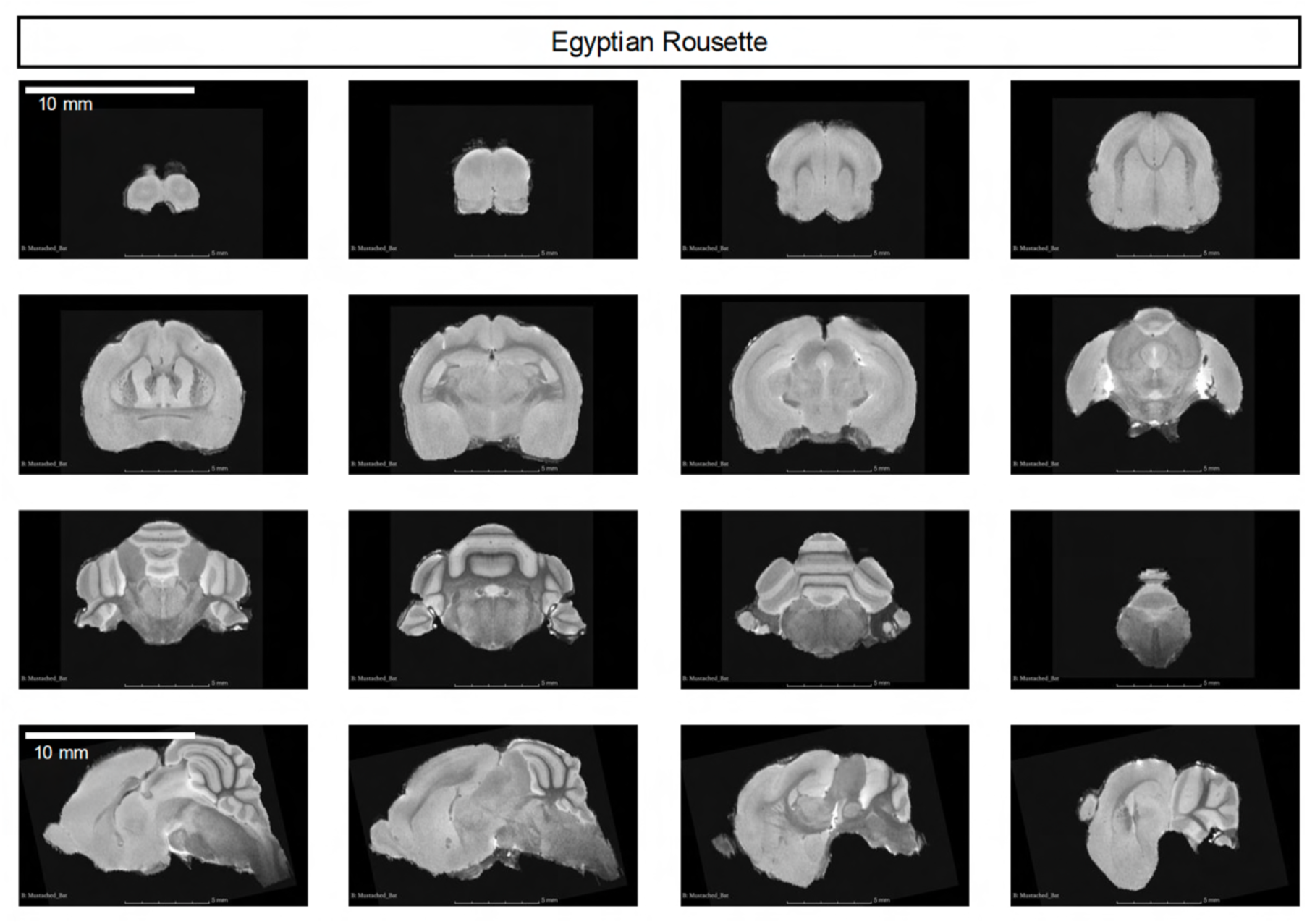
Serial coronal and sagittal T2-weighted MRI sections of the Egyptian rousette. Representative serial T2-weighted MRI sections of the Egyptian rousette brain are shown to provide an overview of whole-brain morphology. Consecutive coronal sections are shown together with sagittal sections to illustrate the anatomical continuity of the forebrain, midbrain, cerebellum, and brainstem. Scale bars, 10 mm.

**Fig. S18.**
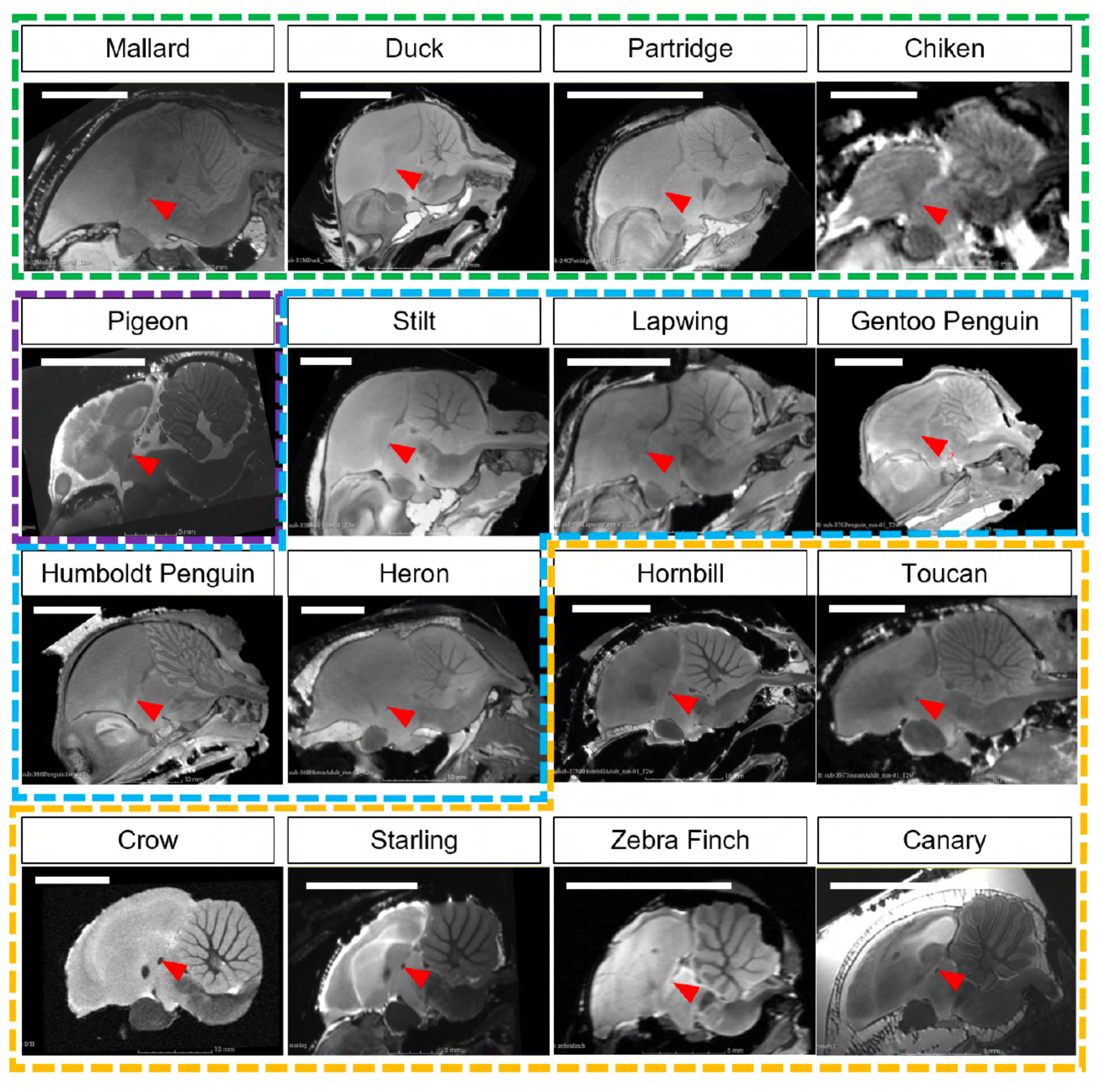
Sagittal T2-weighted MRI sections showing the anterior commissure across bird species. Representative sagittal T2-weighted MRI sections used for the analysis shown in Fig. 2A are shown for the examined bird species. The anterior commissure is indicated by red arrowheads. Species are grouped according to the categories used in the main figure. Scale bars, 10 mm.

**Fig. S19.**
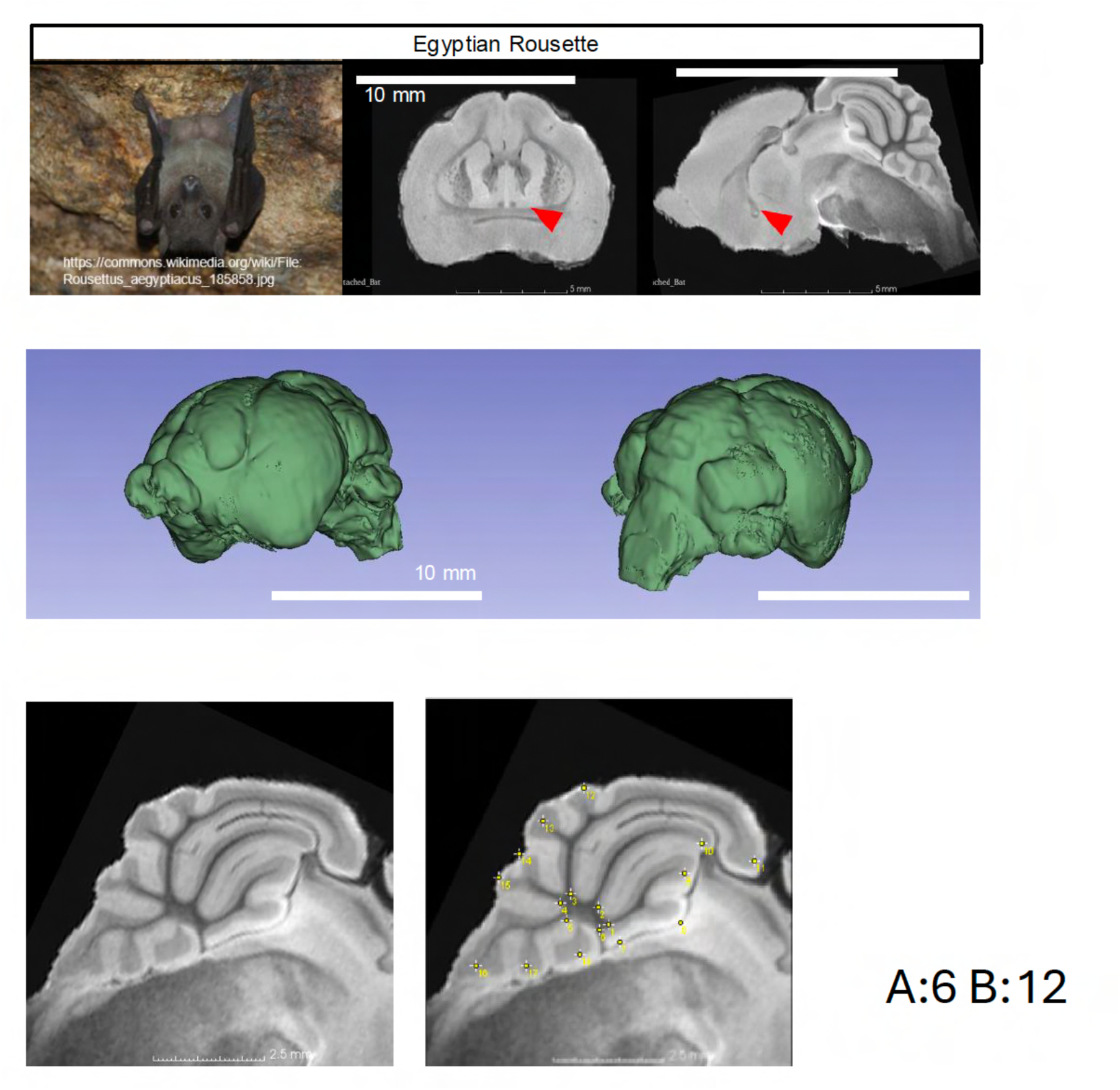
T2-weighted MRI, three-dimensional reconstruction, and cerebellar foliation analysis of the Egyptian rousette. Representative T2-weighted MRI sections and three-dimensional surface reconstructions of the Egyptian rousette brain are shown. The anterior commissure is indicated by red arrowheads in the MRI sections. Cerebellar foliation was assessed using sagittal T2-weighted MRI sections, and representative traces used for branch counting are shown. The number of basal branches and terminal branches is indicated as A and B, respectively. Scale bars, 10 mm.

## Notes

### Competing Interest Statement

The authors have declared no competing interest.

## References

1. E. D. Jarvis, et al., Avian brains and a new understanding of vertebrate brain evolution. Nat. Rev. Neurosci. 6, 151–159 (2005).

2. L. Medina, A. Abellán, Development and evolution of the pallium. Semin. Cell Dev. Biol. 20, 698–711 (2009).

3. S. Olkowicz, et al., Birds have primate-like numbers of neurons in the forebrain. Proc. Natl. Acad. Sci. U. S. A. 113, 7255–7260 (2016).

4. M. Konishi, S. T. Emlen, R. E. Ricklefs, J. C. Wingfield, Contributions of bird studies to biology. Science 246, 465–472 (1989).

5. D. R. Wylie, C. Gutiérrez-Ibáñez, A. N. Iwaniuk, Integrating brain, behavior, and phylogeny to understand the evolution of sensory systems in birds. Front. Neurosci. 9, 281 (2015).

6. O. Vincze, C. I. Vágási, P. L. Pap, G. Osváth, A. P. Møller, Brain regions associated with visual cues are important for bird migration. Biol. Lett. 11, 20150678 (2015).

7. J. R. Corfield, et al., Diversity in olfactory bulb size in birds reflects allometry, ecology, and phylogeny. Front. Neuroanat. 9, 102 (2015).

8. E. D. Jarvis, Learned birdsong and the neurobiology of human language. Ann. N. Y. Acad. Sci. 1016, 749–777 (2004).

9. G. De Groof, A. Van der Linden, Love songs, bird brains and diffusion tensor imaging: DTI OF BRAIN PLASTICITY IN SONGBIRDS. NMR Biomed. 23, 873–883 (2010).

10. A. M. Balanoff, et al., Best practices for digitally constructing endocranial casts: examples from birds and their dinosaurian relatives. J. Anat. 229, 173–190 (2016).

11. C. Poirier, et al., A three-dimensional MRI atlas of the zebra finch brain in stereotaxic coordinates. Neuroimage 41, 1–6 (2008).

12. G. De Groof, et al., In vivo diffusion tensor imaging (DTI) of brain subdivisions and vocal pathways in songbirds. Neuroimage 29, 754–763 (2006).

13. T. Tsurugizawa, et al., A cross-species brain magnetic resonance imaging and histology database of vertebrates. Sci. Data 12, 1206 (2025).

14. A. Van der Linden, et al., Non invasive in vivo anatomical studies of the oscine brain by high resolution MRI microscopy. J. Neurosci. Methods 81, 45–52 (1998).

15. O. Güntürkün, M. Verhoye, G. De Groof, A. Van der Linden, A 3-dimensional digital atlas of the ascending sensory and the descending motor systems in the pigeon brain. Brain Struct. Funct. 218, 269–281 (2013).

16. G. De Groof, et al., A three-dimensional digital atlas of the starling brain. Brain Struct. Funct. 221, 1899–1909 (2016).

17. S. Letzner, A. Simon, O. Güntürkün, Connectivity and neurochemistry of the commissura anterior of the pigeon (Columba livia): Connectivity and neurochemistry of pigeon AC. J. Comp. Neurol. 524, 343–361 (2016).

18. D. T. Ksepka, et al., Tempo and Pattern of Avian Brain Size Evolution. Curr. Biol. 30, 2026–2036.e3 (2020).

19. C. M. Early, A. N. Iwaniuk, R. C. Ridgely, L. M. Witmer, Endocast structures are reliable proxies for the sizes of corresponding regions of the brain in extant birds. J. Anat. 237, 1162–1176 (2020).

20. K. Kverková, et al., The evolution of brain neuron numbers in amniotes. Proc. Natl. Acad. Sci. U. S. A. 119, e2121624119 (2022).

21. R. Yebga Hot, et al., A novel male Japanese quail structural connectivity atlas using ultra-high field diffusion MRI at 11.7 T. Brain Struct. Funct. 227, 1577–1597 (2022).

22. B. Jeurissen, M. Descoteaux, S. Mori, A. Leemans, Diffusion MRI fiber tractography of the brain. NMR Biomed. 32, e3785 (2019).

23. J. L. Hardie, C. R. Cooney, Sociality, ecology and developmental constraints predict variation in brain size across birds. J. Evol. Biol. 36, 144–155 (2023).

24. T. S. Fristoe, C. A. Botero, Alternative ecological strategies lead to avian brain size bimodality in variable habitats. Nat. Commun. 10, 3818 (2019).

25. K. Shiomi, Possible link between brain size and flight mode in birds: Does soaring ease the energetic limitation of the brain? Evolution 76, 649–657 (2022).

26. O. Vincze, Light enough to travel or wise enough to stay? Brain size evolution and migratory behavior in birds: BRAIN SIZE EVOLUTION AND MIGRATION IN BIRDS. Evolution 70, 2123–2133 (2016).

27. F. Sayol, et al., Environmental variation and the evolution of large brains in birds. Nat. Commun. 7, 13971 (2016).

28. L. Z. Garamszegi, A. P. Møller, J. Erritzøe, Coevolving avian eye size and brain size in relation to prey capture and nocturnality. Proc. Biol. Sci. 269, 961–967 (2002).

29. D. S. M. Samia, A. Pape Møller, D. T. Blumstein, Brain size as a driver of avian escape strategy. Sci. Rep. 5, 11913 (2015).

30. M. J. Burish, H. Y. Kueh, S. S.-H. Wang, Brain architecture and social complexity in modern and ancient birds. Brain Behav. Evol. 63, 107–124 (2004).

31. L. Houle, O. Larouche, R. Cloutier, Exploring larval axolotl brain development: Insights into developmental and functional constraints. Evol. Dev. 28, e70034 (2026).

32. S. Kawabe, S. Matsuda, N. Tsunekawa, H. Endo, Ontogenetic shape change in the chicken brain: Implications for paleontology. PLoS One 10, e0129939 (2015).

33. Z. Zhou, et al., Evaluation of the diffusion MRI white matter tract integrity model using myelin histology and Monte-Carlo simulations. Neuroimage 223, 117313 (2020).

34. P. Mukherjee, R. C. McKinstry, Diffusion tensor imaging and tractography of human brain development. Neuroimaging Clin. N. Am. 16, 19–43, vii (2006).

35. T. Nomura, H. Gotoh, K. Ono, Changes in the regulation of cortical neurogenesis contribute to encephalization during amniote brain evolution. Nat. Commun. 4, 2206 (2013).

36. A. Cárdenas, V. Borrell, Molecular and cellular evolution of corticogenesis in amniotes. Cell. Mol. Life Sci. 77, 1435–1460 (2020).

